# Non-classical NMDA receptor subunit GluN3A in a specialized hippocampal region regulates stress-coping strategies in mice

**DOI:** 10.1101/2024.04.24.590964

**Authors:** Wei Zhang, Mingyue Zhao, Jiajie Dai, Linan Zhuo, Haimou Ye, Jiesi Wang, Weiwen Wang

## Abstract

The hippocampus is a highly complex structure, defined by profound structural, functional, and molecular heterogeneity. The GluN3A subunit, which forms non-classic N-methyl-D-aspartate receptors (NMDARs), is enriched in the hippocampal CA1 subregion and has a poorly understood functional role, especially in stress-related behavioral regulation. Here, we employed chronic social defeat stress (CSDS) and GluN3A knockout (KO) mouse models combined with behavioral, molecular, genetic, pharmacological, and circuit-level analyses to investigate the role of hippocampal GluN3A in stress coping. We found that CSDS selectively reduced GluN3A expression in the intermediate CA1 (CA1i) region. Decreased GluN3A expression in CA1i, from either stress or genetic deletion, promoted passive coping behaviors, whereas overexpression of GluN3A in CA1i reversed the phenotype under both conditions. Mechanistically, consistent with GluN3A enrichment in pyramidal neurons, c-fos staining and fiber photometry revealed that GluN3A was essential for CA1i pyramidal neuronal activation during coping behaviors. Further chemogenetic manipulations demonstrated that activating CA1i neurons facilitated active coping, while inhibition induced passivity. Moreover, local administration of D-serine, an agonist of GluN3A-containing NMDARs, rapidly shifted coping behavior from passive to active by activating CA1i neurons via GluN3A-dependent mechanisms. Circuit tracing revealed that CA1i neurons project to multiple sub-hippocampal and cortico-limbic regions, and GluN3A overexpression selectively enhanced downstream neuronal activity in CA3 and infralimbic cortex, but not in the nucleus accumbens. Together, these findings identify GluN3A as a key molecule linking hippocampal subregional specialization with adaptive stress coping, and suggest that the rapid, resilience-promoting effects of D-serine are also GluN3A-dependent.

## INTRODUCTION

The hippocampus is a structurally and functionally heterogeneous structure that plays essential roles in learning, memory, emotional regulation, and adaptive responses to stress[1–5]. It is broadly divided into multiple subfields such as CA1, CA2/3, and the dentate gyrus (DG)[6]. Beyond these classical anatomical partitions, traditional studies have also revealed prominent structural and functional heterogeneity along the longitudinal axis of the hippocampus[3,4,7]. For example, the dorsal hippocampus is primarily involved in spatial learning and cognition, whereas the ventral hippocampus is more associated with emotional regulation and affective behaviors[3,4,8]. Furthermore, advances in molecular profiling, such as spatial transcriptomics, have revealed an even finer organization, showing that the classical dorsal-ventral dichotomy of subfields CA1, CA3, and DG can be refined into dorsal, intermediate, and ventral portions[4,9]. These finer subregions differ in molecular expression, morphology, and projection patterns[9,10]. However, how these distinct neuronal populations and molecular profiles, particularly within the intermediate subregions, execute specific behavioral functions remains largely unclear.

Notably, gene expression profiling has shown a concentration of GluN3A expression in the intermediate and ventral regions of CA1; this specific expression pattern identifies it as a key molecular marker distinguishing the intermediate from the dorsal CA1 subregion [9]. This subunit forms non-classical N-methyl-D-aspartate receptors (NMDARs) with GluN1/N2, exhibiting unique biophysical and pharmacological properties distinct from those of conventional GluN1/GluN2 NMDARs[11,12]. Unlike canonical GluN2-dependent receptors that require glutamate for activation, GluN1/GluN3A assemblies can be directly activated by sensing ambient extracellular glycine or D-serine, generating excitatory glycine receptor-like currents[13–15]. Beyond being glutamate-insensitive, GluN3A-containing receptors also exhibit low Ca^2+^ permeability, significantly reduced Mg^2+^ blockade, and resistance to classical NMDAR antagonists such as D-AP5 and MK-801[13,16–18]. Consistent with these unique features, GluN3A-containing NMDARs have been shown to regulate synaptic maturation and plasticity in several brain regions, including the hippocampus, and have been implicated in multiple affective behavioral phenotypes[16,17,19–22]. Furthermore, growing evidence from preclinical and clinical studies suggests that exogenous D-serine can exert rapid antidepressant-like effects and modulate emotional and cognitive behaviors, underscoring the GluN3A subunit’s potential as a viable target for future clinical translation[23–29]. Collectively, these advances suggest that the subregion-specific expression of GluN3A, combined with the unique properties of the receptors it forms, may represent a key molecular mechanism underlying the hippocampus’s broader spatial and functional heterogeneity; however, its specific hippocampal role remains to be identified, especially in stress-related behavioral regulation.

Chronic stress promotes various negative affective states, including passive coping, anhedonia, and anxiety[8,30–32]. Increasing evidence indicates that chronic stress disrupts hippocampal synaptic transmission and intrinsic excitability, contributing to cognitive deficits, emotional dysregulation, and changes in the stress response [2,32–35]. Furthermore, hippocampal subregions, such as the distinct segments of the CA1, play differential roles in mediating these adverse stress-induced outcomes. For example, the dorsal CA1, as traditionally delineated, is primarily linked to deficits of spatial cognition, working memory, and social function under chronic stress, whereas the ventral CA1 is more closely associated with affective disturbances and anxiety-related behaviors[36–40]. Although the roles of canonical GluN2-containing NMDARs in these processes have been extensively studied[41–43], very little is known regarding the function of non-classical GluN3A-containing NMDARs, which exhibit a distinct spatial expression pattern within the hippocampus, under chronic stress conditions. Recent study has shown that GluN3A deletion in the traditional ventral CA1 region, which encompasses both the CA1 intermediate (CA1i) and CA1 ventral (CA1v) areas, induces anxiety-like behavior and that these excitatory glycine receptors (eGlyR) contribute to corticosterone-dependent modulation of neuronal plasticity, including long-term potentiation[15]. These findings hint that GluN3A may influence stress-related behavioral outcomes and contribute to the functional heterogeneity of CA1. However, GluN3A serving as a molecular marker distinguishing the CA1 dorsal (CA1d) and CA1i subregions[9], it remains unclear which specific hippocampal CA1 loci and behavioral domains GluN3A controls, and how it functionally contributes to stress regulation, particularly under chronic stress exposure.

To address these critical questions, we employed chronic social defeat stress (CSDS) and GluN3A knockout (KO) mouse models, combining comprehensive approaches encompassing behavior, molecular biology, genetic manipulation, microscopic imaging, fiber photometry, pharmacology, and single-cell transcriptomic analysis. This integrative strategy allowed us to delineate the specific contribution of hippocampal GluN3A to passive coping behavior, clarify its role relative to anhedonia, social interaction and anxiety, and identify the neural mechanisms and specific loci underlying this regulation. Our findings provide mechanistic insight into how non-classical NMDARs contribute to stress adaptation, revealing a previously unrecognized role of hippocampal GluN3A in shaping coping strategies under adverse conditions.

## RESULTS

### Effects of CSDS and GluN3A KO on stress-related behaviors and GluN3A expression in the hippocampus

We systematically evaluated various stress-related behaviors induced by CSDS in mice (Fig. 1a, b). Regarding the anhedonia indicator, CSDS mice showed a significant reduction in social interaction ratio (*t_16_* = 5.802, *\*\*\*P*<0.001, Fig. 1c, d) in the social interaction test (SIT) but no significant change in sucrose preference ratio (*P* > 0.05, Supplementary Fig. 1a) in the sucrose preference test (SPT) compared to the control group. In terms of coping behaviors in response to inescapable stress, CSDS significantly increased the immobile time of mice in the tail suspension test (TST) (*t_14_* = 2.877, *\*P* = 0.012, Fig. 1e) and forced swim test (FST) (*t_11_* = 2.240, *\*P* = 0.047, Fig. 1f). In locomotor activity, compared to the control group, CSDS did not affect the total distance traveled by mice (*P*>0.05, Fig. 1g, h). Regarding anxiety-related indicators, CSDS markedly induced anxiety-like behavior in mice, evidenced by a decrease in the number of entries of the central area (*t_13_* = 2.801, *P* = 0.015, Fig. 1i) and the time spent in the central area (Welch-corrected *t_9.157_* = 3.290, *\*P* = 0.009) in the open field test (OFT) (Fig. 1g, j). Additionally, in the elevated plus maze (EPM), CSDS mice exhibited a reduction in the number of open arm entries (Welch-corrected *t_11.43_* = 2.232, *\*P* = 0.047, Fig. 1k, l) and the open arm time ratio (Welch-corrected *t_8.989_* = 2.206, *P* = 0.055, Fig. 1k, m). These results indicated that CSDS successfully induced various negative affective states in mice.

**Figure 1.**
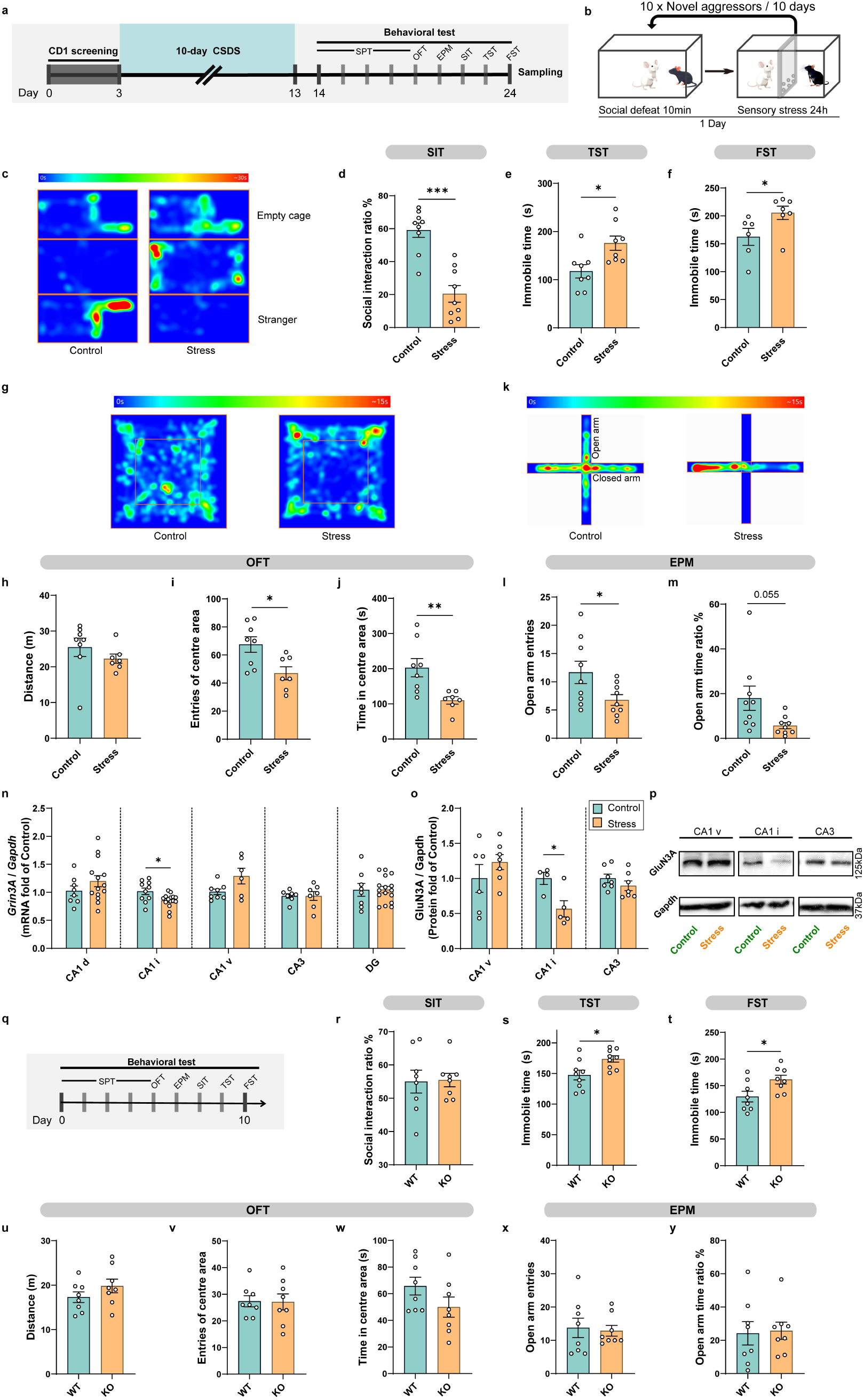
CSDS induces multiple stress-related behaviors and specifically reduces GluN3A expression in the CA1i region, while GluN3A KO specifically promotes passive coping. **a** Experimental scheme of CSDS modeling and behavioral tests. **b** Schematic of CSDS. **c** Example trajectory heatmap of SIT in control mice (left) and stress mice (right). **d** Social interaction ratio of SIT (*n* = 9 per group). **e** Immobility time in the TST (*n* = 8 per group). **f** Immobility time in the FST (*n =* 6 in Control, *n* = 7 in Stress). **g** Example trajectory heatmap of OFT in control mice (left) and stress mice (right). **h** The total distance traveled in OFT (*n =* 8 in Control, *n* = 7 in Stress). **i** The number of entries to centre area of OFT. **j** The time in the centre area of OFT. k Example trajectory heatmap of EPM in control mice (left) and stress mice (right). l The number of entries to open arms of EPM (*n* = 9 per group). m Percentage of time spent in the open arms of EPM. **n** Quantitative data of *Grin3A* mRNA in target brain area from control and stress mice (CA1 d: *n =* 8 in Control, *n* = 14 in Stress; CA1 i: *n =* 10 in Control, *n* = 15 in Stress; CA1 v: *n =* 8 in Control, *n* = 6 in Stress; CA3: *n =* 7 per group; DG: *n =* 8 in Control, *n* = 16 in Stress). **o** Quantitative data of GluN3A protein in target brain area from control and stress mice (CA1 v: *n =* 6 in Control, *n* = 7 in Stress; CA1 i: *n =* 4 in Control, *n* = 5 in Stress; CA3: *n =* 7 per group). **p** Representative western blots for dissected target brain area. **q** Experimental scheme of behavioral studies. **r** Social interaction ratio of SIT (*n* = 8 per group). **s** Immobility time in the TST (*n* = 9 per group). t Immobility time in the FST (*n* = 8 per group). **u** The total distance traveled in OFT (*n* = 8 per group). **v** The entries of centre area in OFT. **w** The time in the centre area of OFT. x The number of entries to open arms of EPM (*n* = 8 per group). **y** Percentage of time spent in the open arms of EPM. *\* P* < 0.05, *\*\*\* P* < 0.001.

Subsequently, we examined the changes in GluN3A expression levels in various emotion-related brain regions, particularly the hippocampus, in mice following CSDS (Supplementary Fig. 1b, c). The quantitative real-time PCR (qRT-PCR) results showed a significant reduction in GluN3A expression in the CA1i region of the hippocampus in CSDS mice compared to the control group (*t_23_* = 2.692, *\*P* = 0.013, Fig. 1n), while there were no significant changes in the other subregions of the hippocampus, including CA1d, CA1v, CA3, and DG (*P* > 0.05, Fig. 1n). The western blot results similarly demonstrated a significant decrease of GluN3A protein expression in the CA1i in CSDS mice (*t_9_* = 2.866, *\*P* = 0.024), with no significant changes observed in CA1v and CA3 (*P* > 0.05, Fig. 1o, p). These data indicated that CSDS downregulated GluN3A expression in the hippocampal CA1i region in mice.

To further examine the relationship between GluN3A and emotion-related behaviors, we conducted behavioral tests on GluN3A KO mice under physiological conditions (Fig. 1q). On the anhedonia indicators, GluN3A KO mice did not exhibit an abnormal ratio of social interactions (*P*>0.05, Fig. 1r) and sucrose preference (*P* > 0.05, Supplementary Fig. 2a) compared to wild-type (WT) mice. In contrast, the results showed that GluN3A KO mice exhibited a passive coping phenotype compared to WT mice, as evidenced by significantly increased immobile time in TST (*t_16_* = 2.778, *\*P* = 0.013, Fig. 1s) and FST (*t_14_* = 2.457, *\*P* = 0.028, Fig. 1t). In addition, GluN3A KO did not influence the locomotor activity (*P*>0.05, Fig. 1u) and anxiety-like behavior (*P*>0.05, Fig. 1v-y).

To ascertain whether GluN3A KO is associated with an increased susceptibility to social defeat stress (SDS), a systematic assessment was conducted to evaluate the effects of subthreshold SDS on emotion-related behaviors in GluN3A KO mice. The results showed GluN3A KO mice subjected to subthreshold SDS exhibited a similar significant increase in immobile time in the FST (Welch-corrected *t_2.520_* = 9.677, *\*P* = 0.031, Supplementary Fig. 2d). Still, no substantial changes observed in social interest, sucrose preference, locomotor activity, and anxiety-like behavior (*P* > 0.05, Supplementary Fig. 2b, c, e, g-i), except a significant reduction in the number of entries of central area in the OFT (*t_13_* = 2.815, *\*P* = 0.015, Supplementary Fig. 2f). The data indicated that the GluN3A KO does not influence the susceptibility of mice to SDS.

Taken together, these results suggested that the decreased expression of GluN3A in the brain was involved in the manifestation of passive coping behaviors, and that the hippocampal CA1i may be a potential brain region for the regulation of stress-coping behaviors by GluN3A.

### Overexpression of the GluN3A decline in CA1i neurons reversed passive coping behaviors

We further investigated the regulatory role of CA1i GluN3A in emotion-related behaviors by using Adeno-associated virus (AAV) to overexpress recombinant GluN3A in the CA1i of CSDS mice (Fig. 2a, b). We found that overexpression of exogenous GluN3A did not significantly affect anhedonia behaviors in CSDS mice (SIT, Fig. 2c; SPT, Supplementary Fig. 3a). In contrast, CA1i-targeted GluN3A overexpression markedly reversed the passive coping phenotype induced by CSDS. Specifically, two-way ANOVA for TST showed a significant interaction effect (*F* _(1, 28)_ = 5.568, *\*P* = 0.026, Fig. 2d), with post-hoc analysis revealing a significant increase in immobile time in the stress: AAV-ZsGreen group compared to the control: AAV-ZsGreen group (False discovery rate-corrected *\*P* = 0.024), while the stress: AAV-Grin3A group showed a significant decrease in immobile time compared to the stress: AAV-ZsGreen group (FDR-corrected *\*P* = 0.016). Two-way ANOVA for FST showed significant main effects of stress (*F* _(1, 28)_ = 5.010, *\*P* = 0.033, Fig. 2e) and viral treatment (i.e., AAV-ZsGreen or AAV-Grin3A) (*F* _(1, 28)_ = 12.99, *\*\*P* = 0.0012) on the immobile time, with post-hoc analysis showing a significant increase in immobile time in the stress: AAV-ZsGreen group compared to the control: AAV-ZsGreen group (FDR-corrected *\*P* = 0.026) and a significant decrease in immobile time in both the control: AAV-Grin3A group (FDR-corrected *\*P* = 0.036) and stress: AAV-Grin3A group (FDR-corrected *\*P* = 0.016) compared to their respective AAV-ZsGreen groups. Furthermore, consistent with the previous experiments, CSDS induced the changes in locomotor indicators and multiple anxiety indicators in OFT and EPM (Fig. 2f-j), but these were not affected by the overexpression of the exogenous GluN3A subunit. These results indicated that the CA1i GluN3A modulated CSDS-induced passive coping behaviors in mice.

**Figure 2.**
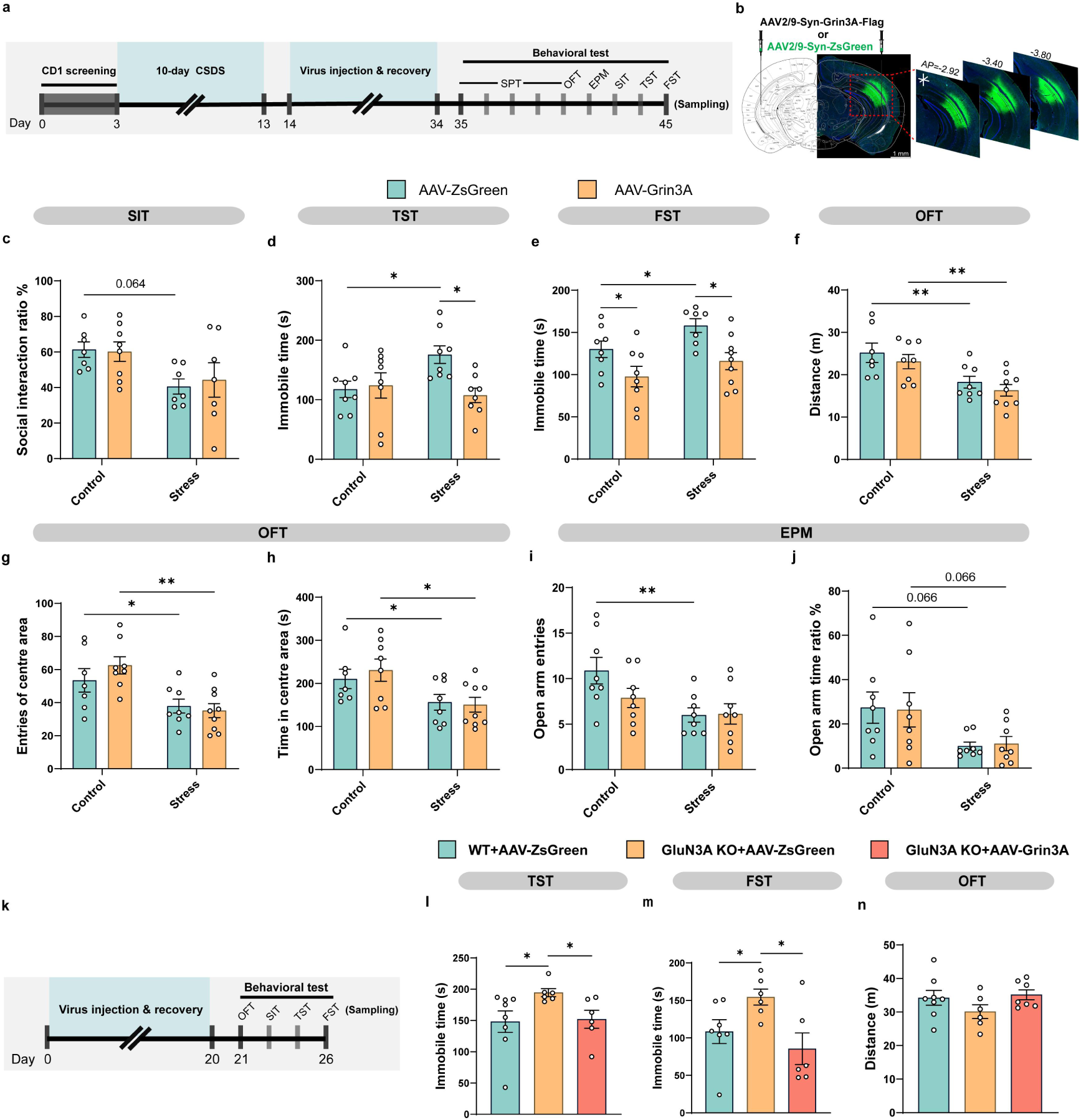
Overexpression of GluN3A in CA1i reverses passive coping behavior in CSDS mice and GluN3A KO mice. **a** Experimental scheme of CSDS modeling, virus injection, and behavior tests. **b** Schematic target locations and immunofluorescence images of ZsGreen or flag expression of Virus in the CA1i. Scale bar, 1 mm. **c** Social interaction ratio of SIT (*n* = 7, 8, 7 and 7 in control:AAV-ZsGreen, control:AAV-Grin3A, stress:AAV-ZsGreen and stress:AAV-Grin3A, respectively). **d** Immobility time in the TST (*n* = 8 per group). **e** Immobility time in the FST (*n* = 8, 8, 7 and 9 in control:AAV-ZsGreen, control:AAV-Grin3A, stress:AAV-ZsGreen and stress:AAV-Grin3A, respectively). **f** The total distance traveled in OFT (*n* = 7, 8, 8 and 9 in control:AAV-ZsGreen, control:AAV-Grin3A, stress:AAV-ZsGreen and stress:AAV-Grin3A, respectively). **g** The number of entries to centre area of OFT. **h** The time in the centre area of OFT. **i** The number of entries to open arms of EPM (*n* = 8 per group). **j** Percentage of time spent in the open arms of EPM. **k** Experimental scheme of virus injection and behavior tests. **l** Immobility time in the TST (*n* = 8, 6 and 6 in WT+AAV-ZsGreen, GluN3A KO+AAV-ZsGreen and GluN3A KO+AAV-Grin3A, respectively). **m** Immobility time in the FST (*n* = 7, 6 and 6 in WT+AAV-ZsGreen, GluN3A KO+AAV-ZsGreen and GluN3A KO+AAV-Grin3A, respectively). **n** The total distance traveled in OFT (*n* = 8, 6 and 7 in WT+AAV-ZsGreen, GluN3A KO+AAV-ZsGreen and GluN3A KO+AAV-Grin3A, respectively). *\* P* < 0.05, *\*\* P* < 0.01.

To further confirm the causal relationship between CA1i GluN3A and stress-coping behaviors, we overexpress recombinant GluN3A in the CA1i region of GluN3A KO mice using AAV (Fig. 2k). Consistent with the findings above, GluN3A overexpression normalized passive coping behaviors without affecting social interaction, locomotor activity, or anxiety-like behavior (*P*>0.05, Fig. 2l-n and Supplementary Fig. 3b-d). Specifically, GluN3A KO mice showed a significant increase in immobile time in TST (*t_12_* = 2.212, *\*P* = 0.047, Fig. 2l) and FST (*t_11_* = 2.309, *\*P* = 0.041, Fig. 2m) compared to WT mice, while GluN3A KO: AAV-Grin3A mice exhibited a significant restoration of immobile time to normal levels in both TST (*t_10_* = 2.696, *\*P* = 0.023) and FST (*t_10_* = 2.912, *\*P* = 0.016) compared to GluN3A KO mice. In summary, hippocampal CA1i GluN3A played a crucial role in regulating stress-induced coping strategies.

### Activity of CA1i pyramidal neurons mediated by GluN3A-dependent regulation of stress-coping behaviors

Having established that GluN3A in the hippocampal CA1i region regulates passive coping behavior in CSDS mice, we next sought to identify the neuronal types that might contribute to this effect. Through reanalysis of single-nuclei RNAseq (snRNAseq) data from previously published studies[44], we found that GluN3A is abundantly expressed in excitatory neurons in the hippocampal CA1 region, and its expression pattern changes after repeated social defeat (RSD) (Fig. 3a, b). Moreover, compared to the Control: WT group, GluN3A expression significantly decreased in excitatory neurons of WT subjected to RSD (*\*P* = 0.037, Fig. 3c, d).

**Figure 3.**
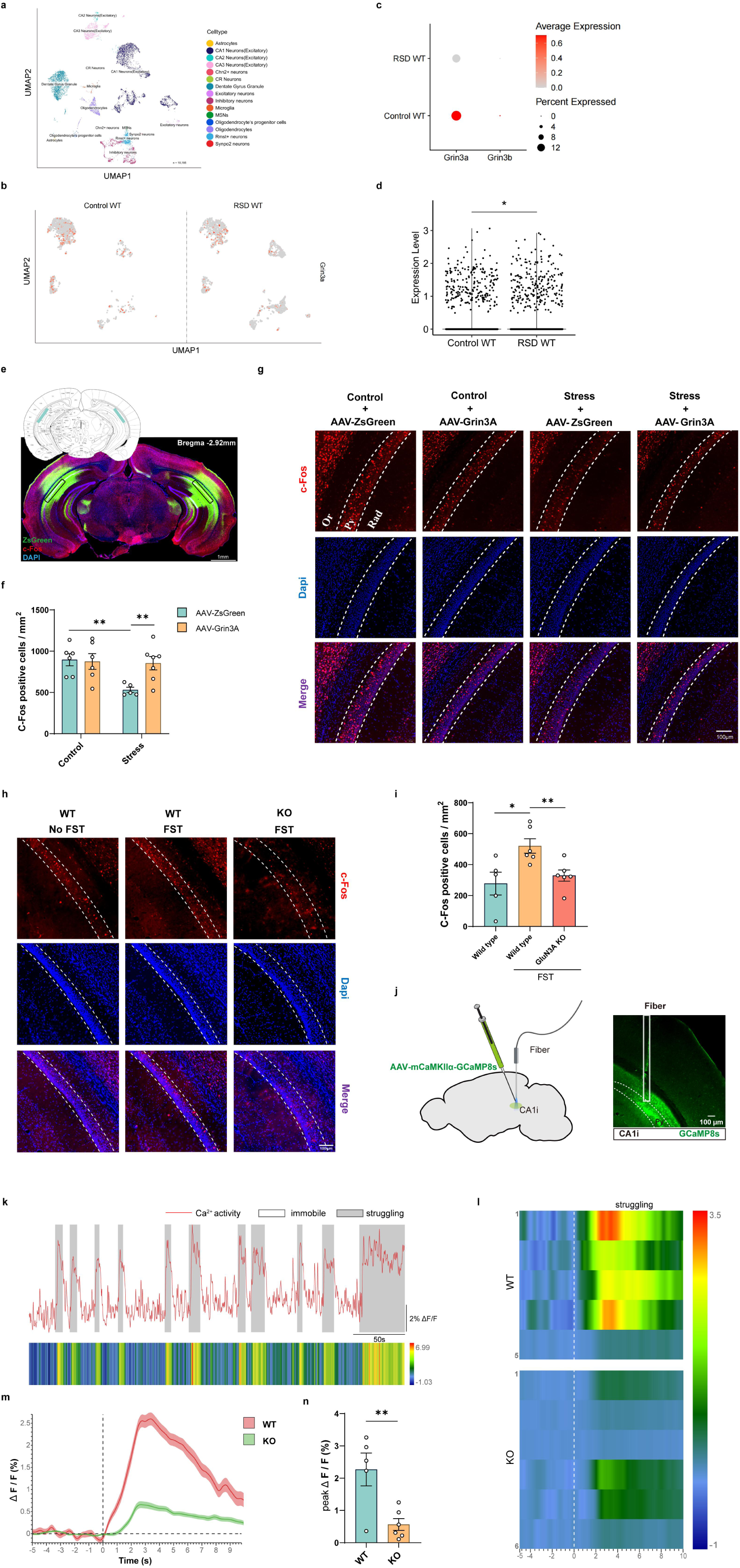
Activity of CA1i pyramidal neurons mediates the GluN3A-dependent regulation of passive coping behavior. **a** Identification of the various cell types in hippocampus using UMAP clustering. **b** Visualization of the distribution of *Grin3A* expression in excitatory neurons in CA1 for each condition: Control WT, RSD WT. **c** Dot plot shows the expression of *Grin3A* in excitatory neurons of each group. The size of the circle representing the percentage of neurons expressing the gene. The color of the circle representing the average level of the gene expression in neurons. **d** Violin plots shows the expression level of *Grin3A* in excitatory neurons of each group. **e** Statistical region of c-Fos-positive cells in CA1i (ZsGreen: green, c-Fos: red, Dapi: blue). Scale bar, 1 mm. **f** Quantification of c-Fos-positive cells in CA1i pyramidal layer neurons of each group of mice. (*n* = 6, 6, 5 and 7 in control:AAV-ZsGreen, control:AAV-Grin3A, stress:AAV-ZsGreen and stress:AAV-Grin3A, respectively, 4 slices for each dot). **g-h** Representative images of c-Fos-positive cells in the CA1i of each group of mice (c-Fos: red, Dapi: blue). Scale bar, 100 μm. **i** Quantification of c-Fos-positive cells in CA1i pyramidal layer neurons of each group of mice. (*n* = 5, 6 and 6 in WT+ No-FST, WT+ FST and GluN3A KO+FST, respectively, 4 slices for each dot). **j** Schematic illustrating fiber placement and representative images of GCaMP8s expression. Scale bar, 100 μm. **k** Top, Example fluorescence trace of the GCaMP signal during TST. Gray shaded area indicates struggling behavior; white area indicates immobile behavior. Bottom, heatmap of fluorescence intensity. **l** Representative heatmaps of GCaMP8s transient ΔF/F event locked to struggling. Each row plots one mice, and a total of 11 mice are illustrated. **m** Dynamics of GCamp8s signaling in mice 5 seconds before and 10 seconds after struggling in the TST (mean ΔF/F ± s.e.m.). **n** Changes in average ΔF/F of WT and KO mice during the struggle (*n =* 5 in WT, *n* = 6 in KO). *\* P* < 0.05, *\*\* P* < 0.01.

Therefore, we further investigated the functional activity of CA1i pyramidal neurons during stress-coping behaviors in the CSDS model under conditions of GluN3A modulation. The results of c-Fos immunostaining showed reduced neuron activity of the pyramidal layer in the CA1i region after the FST in stressed mice compared to controls (Fig. 3e-g), which was restored by overexpressing recombinant GluN3A. Specifically, two-way ANOVA analysis of the number of c-Fos-positive cells showed significant interaction (*F* _(1, 20)_ = 4.820, *\*P* = 0.040, Fig. 3f) and stress main effects (*F* _(1, 20)_ = 6.015, *\*P* = 0.024), with a marginal significant effect for viral treatment (*F* _(1, 20)_ = 3.679, *P* = 0.070). Post-hoc analysis revealed a significant decrease in the number of c-Fos-positive cells in the stress: AAV-ZsGreen group compared to the control: AAV-ZsGreen group (FDR-corrected *\*\*P* = 0.009), while the stress: AAV-Grin3A group showed a significant increase in the number of c-Fos-positive cells compared to the stress: AAV-ZsGreen group (FDR-corrected *\*\*P* = 0.009).

Furthermore, we examined the direct relationship between the GluN3A subunit and the functional activity of CA1i pyramidal neurons after the FST in GluN3A KO mice. Immunostaining results showed that the number of c-Fos-positive cells was significantly increased after FST compared with the basal condition (no FST experience) in WT mice (*t_9_* = 2.884, *\*P* = 0.018), and such activation did not occur in FST-experienced GluN3A KO mice (*t_10_* = 3.232, *\*\*P* = 0.009, Fig. 3h, i), indicating that the activation of pyramidal neurons in the CA1i region was involved in FST behavior, while the absence of GluN3A resulted in this kind of functional defect.

Moreover, we validated the necessity of GluN3A in the activation of excitatory neurons in the CA1i during the TST using fiber photometry (Fig. 3j). During TST, the excitatory-specific GCaMP signal, which characterizes calcium levels, showed a significant correlation with the active state of the mice. The rise of signal was behind the transition from immobile state to struggling state (Fig. 3k). Subsequently, we observed that the average peak amplitude of neuronal responses significantly increased during events when mice transitioned from immobility to struggling (Fig. 3l, m), with GluN3A KO mice showing significantly lower struggle related neuronal activation compared to WT mice (*t_9_* = 3.398, *\*\*P* = 0.008, Fig. 3n).

Collectively, these results indicated that the activity of CA1i pyramidal neurons mediated the GluN3A-dependent regulation of stress-coping behavior in mice.

### Bidirectional chemogenetic modulation of CA1i pyramidal neurons directly mimicked or reversed the stress-induced passive coping behaviors

To directly test the causal role of CA1i pyramidal neuron activity in stress-coping behavior, we employed bidirectional chemogenetic manipulation. We first increased the activity of CA1i neurons using AAV-hM3D(Gq) in combination with different concentrations of clozapine-N-oxide (CNO) (Fig. 4a). The results showed that saline, 0.05 mg/kg, 0.2 mg/kg, and 0.5 mg/kg CNO did not affect the locomotor activity and anxiety-like behavior of normal mice in the OFT and rotarod test (RT) (*P*>0.05, Supplementary Fig. 4a-d). However, one-way ANOVA showed that concentrations of 0.2 mg/kg (FDR-corrected *\*\*P* = 0.009, Fig. 4b-c, Supplementary Fig. 4e) and 0.5 mg/kg (FDR-corrected *\*\*P* = 0.009) CNO effectively increased the number of c-Fos-positive cells at the injection site (*F* _(2, 3)_ = 16.89, *\*P* = 0.023). Therefore, we chose a concentration of 0.2 mg/kg CNO for subsequent chemogenetic activation experiments (Fig. 4d). The results indicated that the activation of CA1i pyramidal neuronal activity in CSDS mice partially enhanced social interest in the SIT (Fig. 4e), and reversed passive coping behavior induced by CSDS in the TST and FST (Fig. 4f, g), but did not alter their locomotor activity and anxiety-like behavior in the OFT (Fig. 4h-j). Specifically, two-way ANOVA for the social interaction ratio showed a significant viral treatment main effect (*F* _(1, 20)_ = 6.983, *\*P* = 0.016, Fig. 4e). Two-way ANOVA for immobile time in the TST showed a significant interaction effect (*F* _(1, 22)_ = 6.923, *\*P* = 0.015, Fig. 4f), with post-hoc analysis showing a significant increase in immobile time in the stress: AAV-mCherry group compared to the control: AAV-mCherry group (FDR-corrected *\*P* = 0.011), while the stress: AAV-hM3D(Gq) group exhibited a significant decrease in immobile time to normal levels compared to the stress: AAV-mCherry group (FDR-corrected *\*P* = 0.011). Two-way ANOVA for immobile time in the FST showed significant stress (*F* _(1, 21)_ = 5.350, *\*P* = 0.031, Fig. 4g) and viral treatment main effects (*F* _(1, 21)_ = 12.34, *\*\*P* = 0.002), with post-hoc analysis showing a significant increase in immobile time in the stress: AAV-mCherry group compared to the control: AAV-mCherry group (FDR-corrected *\*P* = 0.023), and a significant decrease in immobile time in both the control: AAV-hM3D(Gq) group (FDR-corrected *\*P* = 0.026) and stress: AAV-hM3D(Gq) group (FDR-corrected *\*P* = 0.009) compared to their respective AAV-mCherry groups. These results indicated that increased excitatory activity in CA1i neurons can alleviate CSDS-induced passive coping behavior.

**Figure 4.**
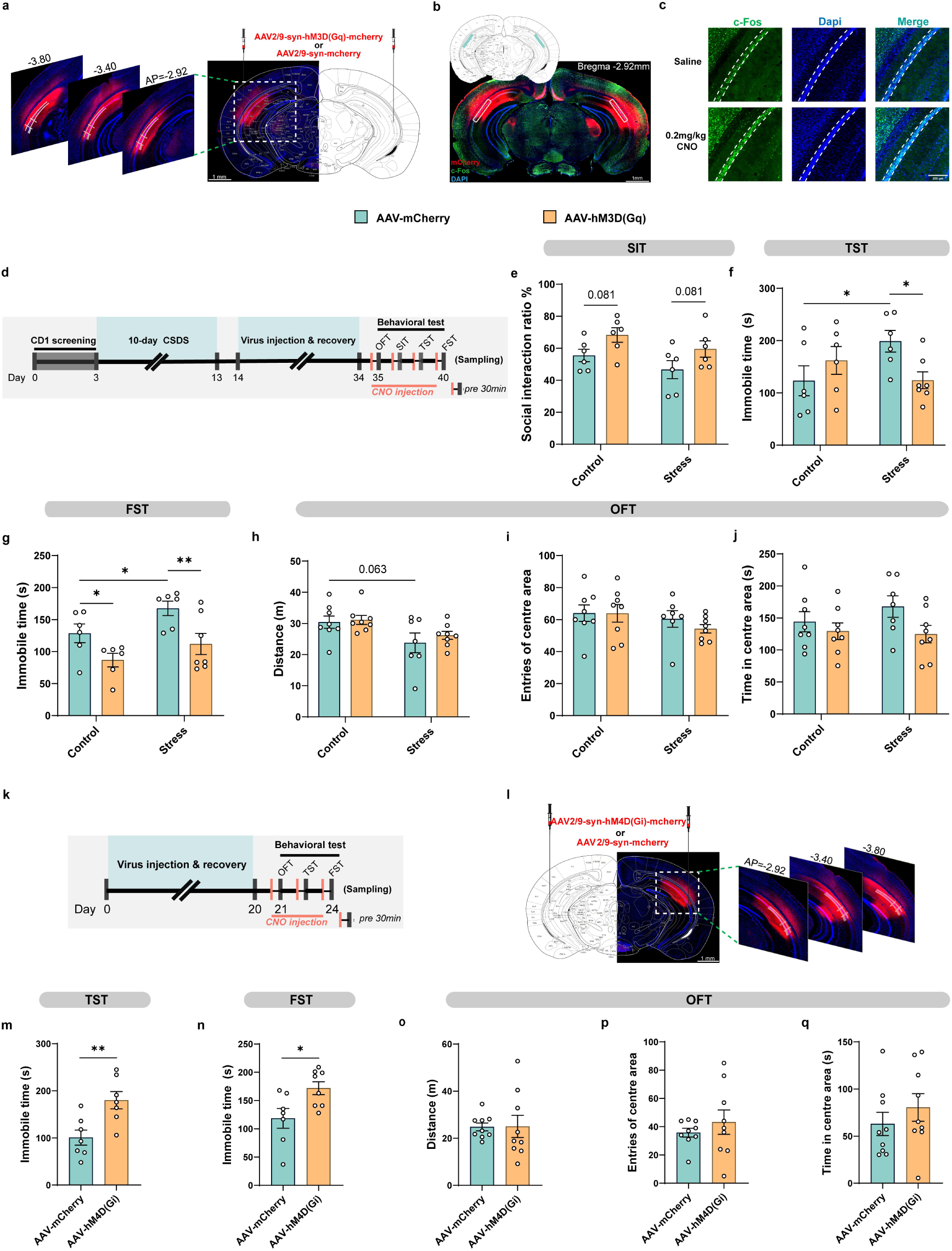
Bidirectional chemogenetic modulation of CA1i pyramidal neurons directly mimics or reverses CSDS-induced passive coping behavior. **a** Schematic target locations and immunofluorescence images of hM3D(Gq)-mCherry or mCherry expression of Virus in the CA1i. Scale bar, 1 mm. **b** Statistical regions of c-Fos-positive cells in CA1i (mCherry: red, c-Fos: green, Dapi: blue). Scale bar, 1 mm. **c** Representative images of c-Fos-positive cells in the CA1i of the saline and 0.2 mg/kg CNO treatment mice (c-Fos: green, Dapi: blue). Scale bar, 200 μm. **d** Experimental scheme of CSDS modeling, virus injection, and behavior tests without or with chemogenetic activation. **e** Social interaction ratio of SIT (*n* = 6 per group). **f** Immobility time in the TST (*n* = 6, 6, 6 and 7 in control:AAV-mCherry, control:AAV-hM3D(Gq), stress:AAV-mCherry and stress:AAV-hM3D(Gq), respectively). **g** Immobility time in the FST (*n* = 6, 6, 6 and 7 in control:AAV-mCherry, control:AAV-hM3D(Gq), stress:AAV-mCherry and stress:AAV-hM3D(Gq), respectively). **h** The total distance traveled in OFT (*n* = 8, 8, 7 and 8 in control:AAV-mCherry, control:AAV-hM3D(Gq), stress:AAV-mCherry and stress: AAV-hM3D(Gq), respectively). **i** The number of entries to centre area of OFT. **j** The time in the centre area of OFT. **k** Experimental scheme of virus injection, and behavior tests without or with chemogenetic inactivation. **l** Schematic target locations and immunofluorescence images of hM4D(Gi)-mCherry or mCherry expression of Virus in the CA1i. Scale bar, 1 mm. **m** Immobility time in the TST (*n* = 7 per group). **n** Immobility time in the FST (*n =* 7 in AAV-mCherry, *n* = 8 in AAV-hM4D(Gi)). **o** The total distance traveled in OFT (*n* = 9 per group). **p** The number of entries to centre area of OFT. **q** The time in the centre area of OFT. *\* P* < 0.05, *\*\* P* < 0.01.

On the other hand, we inhibited the activity of CA1i pyramidal neurons in normal mice using AAV-hM4D(Gi) (Fig. 4k, l). We found that reducing the activity of the neurons significantly increased immobile time in the TST (*t_12_* = 3.261, *\*\*P* = 0.007, Fig. 4m) and FST (*t_13_* = 2.622, *\*P* = 0.021, Fig. 4n), but did not induce abnormalities in locomotor activity and anxiety-like behavior in the OFT (*P*>0.05, Fig. 4o-q), indicating that the inhibition of neuronal activity in CA1i can directly induce passive coping phenotype.

Overall, our findings suggest that the activity of CA1i pyramidal neurons may underlie GluN3A’s regulation of stress-coping behavior.

### Potentiation of GluN3A-mediated CA1i pyramidal neuron activity was crucial for the active coping strategies induced by D-serine

We further explored whether GluN3A participates in the rapid D-serine–induced shift of stress-coping strategies within the hippocampus. By injecting D-serine into the bilateral CA1i region of CSDS mice and GluN3A KO mice before behavioral testing (Fig. 5a, b), we found that D-serine could significantly reverse the passive coping behavior induced by CSDS (Fig. 5c, d), while it had no significant effect on locomotor activity and anxiety-like behavior in the OFT in all groups (*P*>0.05, Fig. 5e-g). Specifically, two-way ANOVA for immobile time in the TST and FST showed significant interaction (Control-cerebrospinal fluid (CSF), Control-D-serine, Stress-CSF and Stress-D-serine group, TST: *F* _(1, 27)_ = 6.084, *\*P* = 0.020; FST: *F* _(1, 27)_ = 5.106, *\*P* = 0.032, Fig. 5c, d) and drug treatment main effects (TST: *F* _(1, 27)_ = 11.60, *\*\*P* = 0.002; FST: *F* _(1, 27)_ = 11.19, *\*\*P* = 0.002), with post-hoc analysis showing a significant increase in immobile time in the stress: CSF group compared to the control: CSF group in both tests (TST: FDR-corrected *\*P* = 0.017; FST: FDR-corrected *\*P* = 0.032), while the stress: D-serine group exhibited a significant decrease in immobile time compared to the stress: CSF group (TST: FDR-corrected *\*\*\*P* < 0.001; FST: FDR-corrected *\*\*P* = 0.002). Interestingly, compared to the control (WT):D-serine group, the GluN3A KO:D-serine group still exhibited significantly increased immobile time in both the TST (*t_13_* = 2.470, *^+^P* = 0.028) and FST (*t_13_* = 3.225, *^++^P* = 0.007), indicating that GluN3A is required for D-serine–induced behavioral improvement. Additionally, simultaneously inhibiting the activity of CA1i neurons in CSDS mice while administering D-serine, we found that the stress-hM4D(Gi) group showed a significant increase in immobile time in both the TST (*t_12_* = 4.628, *^###^P* < 0.001) and FST (*t_12_* = 2.671, *^#^P* = 0.020) compared to the Stress-mCherry group, suggesting that suppressing CA1i pyramidal neuronal activity blocked the active coping effect of D-serine.

**Figure 5.**
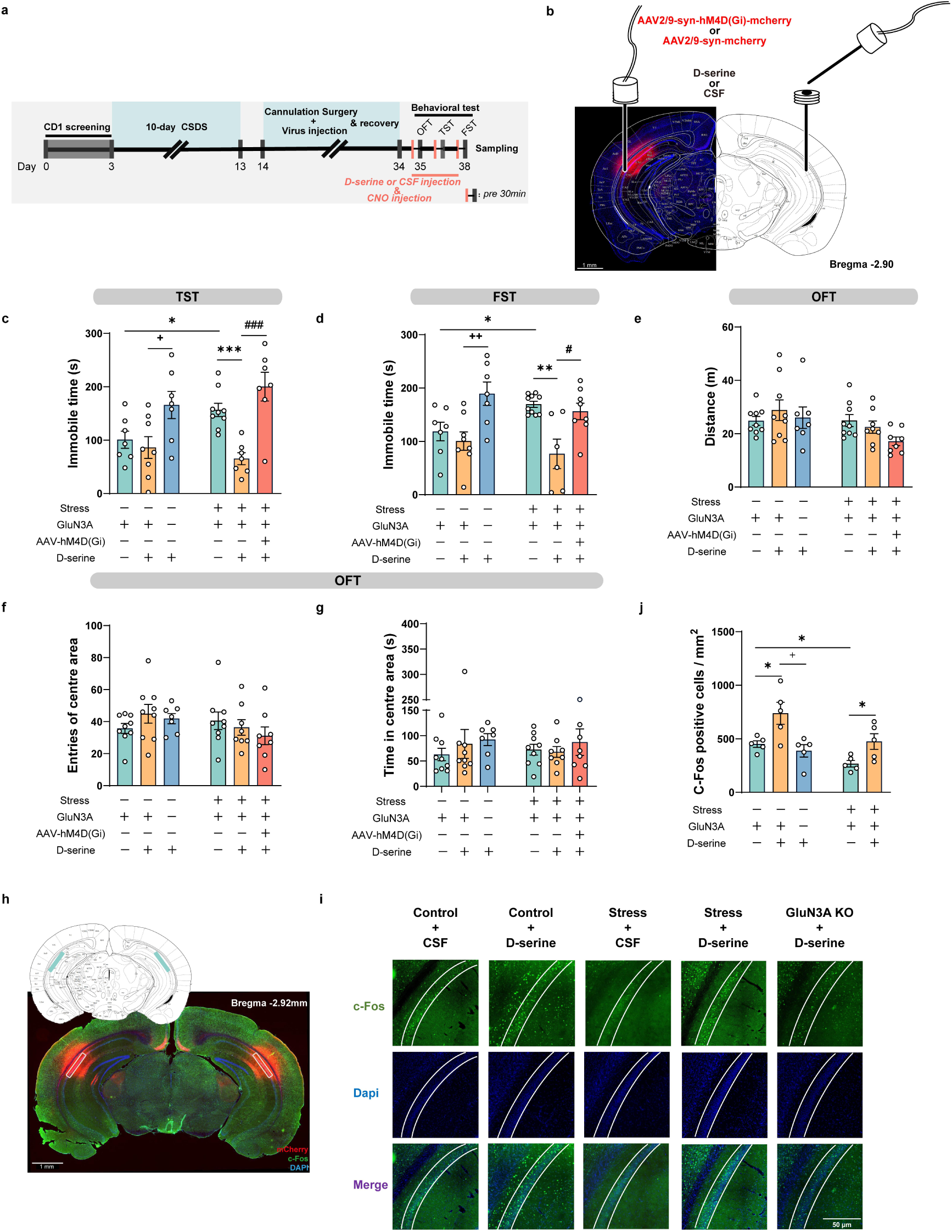
GluN3A-dependent potentiation of CA1i pyramidal neuronal activity is essential for the rapid shift in coping strategy by D-serine. **a** Experimental scheme of CSDS modeling, virus injection, cannula implantation, and behavior tests without or with D-serine delivery. **b** Schematic target locations injection of D-serine (CSF) and Virus in the CA1i. Scale bar, 1 mm. **c** Immobility time in the TST (*n* = 7, 8, 7, 9, 7 and 7 in Control-CSF, Control-D-serine, GluN3A KO:D-serine, Stress-CSF, Stress-D-serine and stress-D-serine-hM4D(Gi), respectively). **d** Immobility time in the FST (*n* = 7, 8, 7, 10, 6 and 8 in Control-CSF, Control-D-serine, GluN3A KO:D-serine, Stress-CSF, Stress-D-serine and stress-D-serine-hM4D(Gi), respectively). **e** The total distance traveled in OFT (*n* = 9, 9, 7, 9, 8 and 8 in Control-CSF, Control-D-serine, GluN3A KO:D-serine, Stress-CSF, Stress-D-serine and stress-D-serine-hM4D(Gi), respectively). **f** The number of entries to centre area of OFT. **g** The time in the centre area of OFT. **h** Statistical regions of c-Fos-positive cells in CA1i (mCherry: red, c-Fos: green, Dapi: blue). Scale bar, 1 mm. **i** Representative images of c-Fos-positive cells in the CA1i of each group of mice (c-Fos: green, Dapi: blue). Scale bar, 50 μm. **j** Quantification of c-Fos-positive cells in CA1i pyramidal layer neurons of each group of mice. (per mm, *n* = 5 per group, 4 slices for each dot). *\* P* < 0.05, *\*\* P* < 0.01, *\*\*\* P* < 0.001; *^+^ P* < 0.05, *^++^ P* < 0.01; *^#^ P* < 0.05, *^###^P* < 0.001.

Furthermore, we performed immunostaining for c-Fos in the CA1i pyramidal layer neurons of mice from the groups above after the FST (Fig. 5h). The results showed that D-serine significantly reversed the decrease in the number of c-Fos-positive cells induced by CSDS in the CA1i pyramidal layer (Fig. 5i, j). Specifically, two-way ANOVA for the number of c-Fos-positive cells showed significant stress (Control-CSF, Control-D-serine, Stress-CSF, and Stress-D-serine group, *F* _(1, 16)_ =11.23, *\*\*P* = 0.004, Fig. 5j) and drug treatment effects (*F* _(1, 16)_ =13.78, *\*\*P* = 0.002). Post-hoc analysis showed a significant decrease in the number of c-Fos-positive cells in the stress: CSF group compared to the control: CSF group (FDR-corrected *\*P* = 0.037), while both the control: D-serine group (FDR-corrected *\*P* = 0.016) and stress: D-serine group (FDR-corrected *\*P* = 0.045) showed a significant increase in the number of c-Fos-positive cells compared to their respective CSF groups. However, in GluN3A KO:D-serine mice, the number of c-Fos-positive cells in the CA1i pyramidal layer did not rise to the level of the control (WT):D-serine group (*t_8_* = 2.967, *^+^P* = 0.018). In summary, D-serine facilitates a rapid shift from passive to active coping behavior by potentiating GluN3A-dependent CA1i pyramidal neuron activity, highlighting GluN3A signaling in this hippocampal subregion as a crucial modulator of adaptive stress responses.

### Projections of CA1i neurons and downstream circuits may be involved in GluN3A-mediated regulation of stress-coping behaviors

To identify the downstream targets of hippocampal CA1i neurons potentially involved in GluN3A-mediated behavioral regulation, we first performed projection-mapping analyses using the Allen Brain Atlas database and AAV-mediated tracing experiments. According to the Allen Brain connectivity data, CA1i neurons send prominent projections to several sub-hippocampal and limbic related brain regions, including the CA3, nucleus accumbens (NAc), infralimbic cortex (IL), BLA, and lateral entorhinal cortex (LEC) (Fig. 6a). Consistent with these database results, injection of AAV2/9-syn-ZsGreen into the CA1i region resulted in robust fluorescent projection signals in these downstream areas, particularly in the CA3, NAc core (AcbC) and shell (AcbSh), IL, BLA, and LEC (Fig. 6b), confirming that CA1i pyramidal neurons form extensive efferent connections with emotion-related circuits.

**Figure 6.**
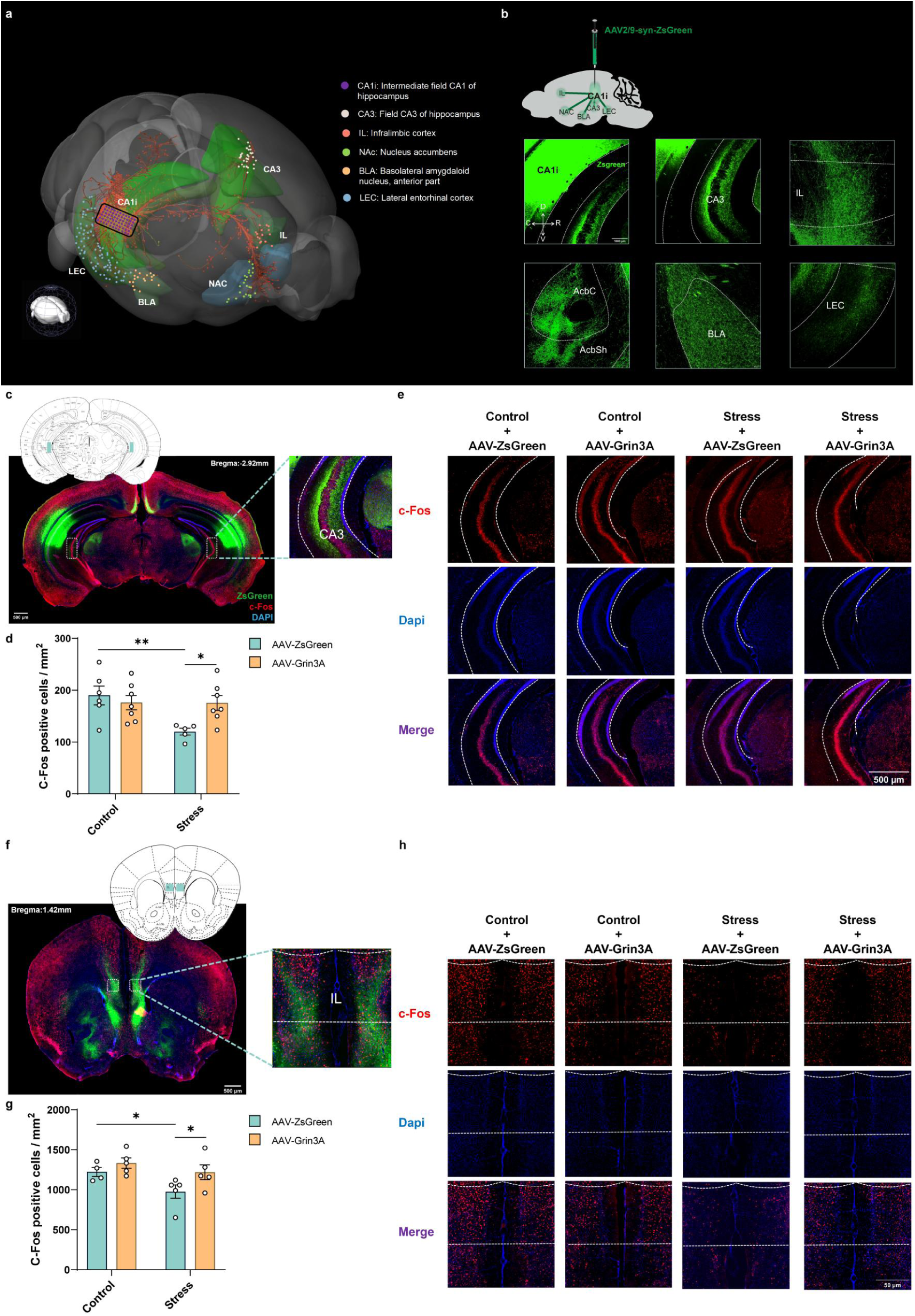
GluN3A-mediated activation of CA1i pyramidal neurons selectively influenced neuronal activity in downstream CA3 and IL circuits. **a** Projection map of the hippocampal CA1i region generated from the Allen Brain Atlas database. **b** Representative images of virus tracing results following AAV2/9-syn-ZsGreen injection into the CA1i region. Scale bar, 1000 μm. **c** Statistical region of c-Fos-positive cells in CA3 (ZsGreen: green, c-Fos: red, DAPI: blue). Scale bar, 500 μm. **d** Quantification of c-Fos-positive cells in CA3 neurons of each group of mice (n = 6, 7, 5 and 7 in control:AAV-ZsGreen, control:AAV-Grin3A, stress:AAV-ZsGreen and stress:AAV-Grin3A, respectively, 4 slices for each dot). **e** Representative images of c-Fos-positive cells in the CA3 of each group of mice (c-Fos: red, Dapi: blue). Scale bar, 500 μm. **f** Statistical region of c-Fos-positive cells in IL (ZsGreen: green, c-Fos: red, DAPI: blue). Scale bar, 500 μm. **g** Quantification of c-Fos-positive cells in IL neurons of each group of mice (n = 4, 5, 5 and 5 in control:AAV-ZsGreen, control:AAV-Grin3A, stress:AAV-ZsGreen and stress:AAV-Grin3A, respectively, 4 slices for each dot). h Representative images of c-Fos-positive cells in the IL of each group of mice (c-Fos: red, Dapi: blue). Scale bar, 50 μm. *\* P* < 0.05, *\*\* P* < 0.01.

Next, we examined whether overexpression of the GluN3A subunit in CA1i neurons could influence neuronal activation in these projection targets following chronic stress. c-Fos immunostaining was performed in representative downstream regions after the FST in control and CSDS mice receiving either AAV-ZsGreen or AAV-Grin3A injections in the CA1i region (Fig. 6c-h, Supplementary Fig. 5a-d). In the CA3 region, a two-way ANOVA revealed a significant stress × viral treatment interaction (*F* _(1, 21)_ = 5.641, *\*P* = 0.027, Fig. 6d) and a significant stress main effect (*F* _(1, 21)_ = 5.816, *\*P* = 0.025). Post hoc analysis showed that in the AAV-ZsGreen group, CSDS significantly decreased the number of c-Fos–positive neurons in the CA3 region compared with controls (FDR-corrected *\*\*P* = 0.009). In contrast, overexpression of GluN3A in CA1i neurons reversed this reduction, restoring c-Fos expression to control levels (FDR-corrected *\*P* = 0.017). In the IL, both stress (*F* _(1, 15)_ = 5.368, *\*P* = 0.035, Fig. 6g) and viral treatment (*F* _(1, 15)_ = 5.182, *\*P* = 0.038) exerted significant main effects. Post hoc analysis indicated that in the AAV-ZsGreen group, CSDS significantly reduced c-Fos–positive cell counts in the IL (FDR-corrected *\*P* = 0.049), whereas GluN3A overexpression markedly increased c-Fos expression levels relative to the stress: AAV-ZsGreen group (FDR-corrected *\*P* = 0.049). In contrast, in both the AcbSh and AcbC regions, there were no significant effects observed (*P*>0.05, Supplementary Fig. 5b, c). These findings suggest that GluN3A modulation in the CA1i primarily affects activity in specific CA3 and IL targets, rather than in the NAC.

Together, these results indicate that CA1i pyramidal neurons establish strong anatomical and functional connections with emotion-related structures, and that GluN3A-dependent regulation of stress-coping behaviors may occur through selective modulation of CA1i-CA3 or CA1i-IL circuit activity rather than global changes across all projection targets.

## DISCUSSION

In the present study, we systematically investigated a previously unrecognized role of the GluN3A-containing non-classical NMDARs within the hippocampal CA1i region in regulating stress-coping strategies. Our results demonstrated that chronic stress induced a specific decrease in GluN3A expression in the hippocampal CA1i region, and that CA1i GluN3A explicitly regulates the passive coping behaviors in the TST and FST. Mechanistically, the effects of GluN3A on stress-coping may be mediated through the activation of CA1i pyramidal neurons. Moreover, we found that D-serine administration facilitated a rapid shift from passive to active coping by activating GluN3A-dependent CA1i neurons. Finally, circuit-level analyses revealed that the selective, GluN3A-mediated activation of CA1i-CA3 and CA1i-IL pathways, but not CA1i-NAc, may serve as a potential circuit substrate for stress adaptation. Collectively, these findings highlight a critical role for GluN3A in response to chronic stress, and its ability to shape adaptive coping through hippocampal CA1i pyramidal neurons and specific projection circuits, and reveal its contribution to the rapid shift in coping strategy induced by D-serine.

First, by focusing on the non-classical NMDAR subunit GluN3A and its distinctive expression pattern in refined hippocampal subregions, we found that mice exhibited a selective reduction in GluN3A expression in the CA1i region, but not in the traditional, more broadly dorsal and ventral subareas, after CSDS. This finding provides functional significance for the Hippocampal Gene Expression Atlas (HGEA) constructed by Bienkowski et al., which divides the CA1 into four subregions: CA1d, CA1i, CA1v, and the ventral tip (CA1vv), and identifies GluN3A as a key molecular marker distinguishing CA1i from CA1d[9]. Moreover, our causal manipulations (both genetic knockout and selective CA1i overexpression in CSDS or KO animals) confirmed that CA1i GluN3A bidirectionally regulates passive coping behaviors in the TST and FST, without affecting social interest or anxiety-like behaviors. This demonstrates a regionally and behaviorally specific regulatory role for CA1i GluN3A-expressing neurons in stress coping. This behavioral specificity for coping refines previous reports. For instance, while one study linked global hippocampal GluN3A influences to both helplessness and anxiety[45], another suggested that GluN3A in the traditional ventral hippocampus modulates anxiety-like behaviors and is essential for neurons’ response to corticosterone[15]. Our results reveal a more precise function, identifying GluN3A as a key molecular determinant of stress response and coping flexibility, and pinpointing CA1i as a critical functional locus linking molecular heterogeneity to behavioral specificity during stress adaptation.

Coping behavior represents a core dimension of the stress response, reflecting the individual’s ability to adjust strategies dynamically when facing adversity[46,47]. Active coping is associated with resilience and positive affect, whereas passive coping (helplessness) is linked to depression vulnerability[48–51]. Clinically, impairments in coping flexibility are a hallmark of major depressive disorder and other stress-related psychopathologies[52–54]. In human studies, genetic variations in the GRIN3A locus have been implicated in bipolar disorder and schizophrenia[55,56], lending clinical relevance to our findings and further suggesting that the CA1i GluN3A-mediated pathway we identified may contribute to the pathophysiology of related mental illnesses. Thus, identifying CA1i GluN3A as a regulator of coping strategies may provide novel therapeutic entry points for disorders characterized by maladaptive stress responses.

Our study further elucidates the neuronal mechanism by which GluN3A modulates stress-coping strategies, suggesting that modulation of CA1i neuronal excitability, primarily in pyramidal neurons, is the key mediator. Several convergent lines of evidence support this conclusion. First, snRNAseq data revealed a significant downregulation of GluN3A expression in CA1 excitatory neurons of CSDS-exposed mice[44], a finding consistent with the identified neuronal hypoactivity observed in CSDS and GluN3A KO mice in the current study. Second, both stress exposure and GluN3A KO decreased CA1i c-Fos expression after coping behaviors, and KO animals further showed attenuated struggling-related calcium transients in pyramidal neurons during TST. Third, overexpression of CA1i GluN3A restored c-fos-represented neuronal activity and rescued passive coping behavior. Critically, Chemogenetic manipulation further established a causal link between CA1i neuron excitability and coping behavior: activation promoted active coping, while inhibition induced passive coping. Although we cannot exclude the potential contribution of GluN3A-expressing inhibitory neurons, our results suggest that pyramidal neurons are the main effector population translating GluN3A signaling into behavioral outcomes.

One plausible mechanism underlying this regulatory role in CA1i neuron activity may involve the unique properties of GluN3A-containing NMDARs, which are inherently glutamate-insensitive but responsive to extracellular glycine or D-serine, enabling CA1i pyramidal neurons to sense glial-derived glycine or D-serine released under stress conditions [13,15,16]. Chronic stress-induced downregulation of GluN3A may disrupt this alternative excitatory pathway, thereby dampening pyramidal neuron excitability and biasing behavioral output toward passive coping. Recent electrophysiological evidence supports the core premise of this hypothesis, which indicates that excitatory glycine receptors are present on ventral hippocampal pyramidal neurons and mediate distinct synaptic dynamics [15]. Furthermore, the proposed link between GluN3A and neuronal excitability is further strengthened by findings in other brain regions, such as the olfactory bulb, where GluN3A KO also reduces c-Fos expression and CaMKII phosphorylation[57]. Finally, the causal link between CA1 pyramidal neuron hypoactivity and passive coping is consistent with previous studies, as inhibition of these neurons has been shown to promote depression-like behaviors, including stress-coping behaviors represented by the FST and TST[34,35]. This convergence of evidence strengthens the hypothesis that this non-classical GluN3A-containing NMDAR pathway modulates CA1i pyramidal neuron excitability.

Our findings on D-serine further underscore the translational significance of GluN3A signaling. As a high-affinity ligand for the glycine-binding site of GluN3 subunits[13,58], D-serine and its derivatives have been reported to exert rapid antidepressant effects in both preclinical and clinical studies, with the hippocampus being a critical site of action[23,28,59]. Given that passive coping is a behavioral endophenotype targeted by antidepressant treatment[60–62], we explored the mechanistic link between D-serine, GluN3A, and stress-coping behaviors. We found that D-serine microinjection into the CA1i rapidly shifted coping behavior from passive to active, accompanied by increased CA1i pyramidal neuron activity. Remarkably, this effect was abolished in GluN3A KO mice or when CA1i neuron activity was suppressed, indicating that GluN3A-dependent excitation of CA1i pyramidal neurons is essential for D-serine’s behavioral efficacy. Given that D-serine itself has also been reported to have extensive links, including endogenous changes and exogenous administration, to symptom or cognitive improvements in clinical and pre-clinical settings[63–66], our findings suggest that the GluN3A-dependent pathway in CA1i is not only critical for adaptive coping but may also offer a promising and mechanistically targetable path for the development of therapeutic interventions or cognitive enhancements in related conditions.

Finally, our initial exploration of CA1i projection analysis points toward potential circuit-level mechanisms to explain the significant role of GluN3A. We demonstrated that CA1i neurons project extensively to sub-hippocampal and limbic-related regions, including CA3, IL, and NAc. Furthermore, our results showed that GluN3A overexpression selectively enhanced downstream neuronal activation in CA3 and IL, but not NAc. These projection circuits, although not specific to CA1i, have previously been reported to be associated with stress-coping strategies of mice [30,67,68]. This alignment with previous reports strongly suggests that CA1i-CA3 and CA1i-IL projections may be the functional circuits underlying GluN3A-dependent regulation of stress coping by CA1i neurons.

In conclusion, our study identifies non-classical GluN1/GluN3A NMDARs in the hippocampal CA1i as key regulators of adaptive stress responses by selectively activating pyramidal neurons and defined projection circuits. While further studies are warranted to determine the exact roles of GluN3A-expressing pyramidal neurons and their downstream circuit targets, our findings bridge hippocampal structural and transcriptional heterogeneity with behavioral specialization, offering new insight into the neural basis of coping and resilience. Furthermore, the discovery that D-serine exerts GluN3A-dependent rapid modulation of coping behavior highlights a translational opportunity for targeting non-classical NMDARs in clinical usage, such as the treatment of depression and the improvement of stress-related symptoms.

## MATERIALS AND METHODS

### Animals

All animal procedures were approved by the Institutional Animal Care and Use Committee of the Institute of Psychology, Chinese Academy of Sciences. C57BL/6N male mice and CD-1 (ICR) male mice were purchased from Charles River, Beijing, China. *Grin3A*^-/-^ (GluN3A KO) mice (C57BL/6J background) were provided by Tifei Yuan’s laboratory. This model was established by the construction of a specific small guide RNA (sgRNA) vector to direct Cas9 to delete exon 2 of the GluN3A gene. The GluN3A KO and WT littermates used in this study were bred from the *Grin3A*^+/-^ mice in the individually ventilated cage (IVC). All mice were housed in a specific-pathogen-free (SPF) environment with a temperature of 22±2℃, relative humidity of 40-70%, and a 12-hour (h) light-dark cycle (7:00 am-19:00 pm). The experimental mice had free access to food and water throughout the experimental period.

### CSDS model

CSDS was performed as previously described[69,70]. Male CD1 mice, singly housed for 10-24 weeks, were screened based on aggression intensity. CD1 mice displaying attacks toward unfamiliar “intruders” within 3 minutes (min) for 3 consecutive days were designated as “residents” for subsequent stress experiments. 3 days before the CSDS experiment, “resident” CD1 mice were placed into the CSDS cage to establish territorial awareness. During each day of the 10-day CSDS period, adult C57BL/6N mice were introduced into the “resident” CD1 mouse home cage and subjected to physical attacks for 10 min. For the remainder of the day, a transparent perforated PVC divider was used to separate them while maintaining visual, auditory, and olfactory contact. For 10 successive days, a C57BL/6N mouse was introduced to a novel aggressive CD1 mouse home cage every day. C57BL/6N control mice were housed in the CSDS cage with a divider to avoid body contact, and a new cage mate was introduced every 24 hours (h) for 10 days. After the social defeat stress, the experimental mice were individually housed until all tests were completed.

### Subthreshold SDS

Experiments were performed with GluN3A KO mice involving three consecutive 5 min physical interactions with a novel CD-1 mice, with a 15 min interval between each interaction[71].

### Behavioral assessments

This study assessed the anhedonia symptom using the SPT (sensory anhedonia) and the three-chamber SIT (social anhedonia). Locomotor activity was evaluated through the OFT and RT, while coping behavior was measured using the TST and FST. Anxiety symptoms were examined through the OFT and EPM.

All behavioral tests were conducted during the light cycle, with at least a 24h interval between different tests. 30min before each behavioral test, experimental mice were placed in a dimly lit testing room to acclimate to the environment. After each mouse completed the behavioral test, the apparatus was cleaned with 70% alcohol to eliminate residual mouse odors and traces. All test procedures were video-recorded.

#### SPT

During adaptation phase, mice were acclimatized for 48h by simultaneously giving one bottle of 1% sucrose solution and one bottle of pure water, and the bottle positions were switched after 24h of acclimatization in order to eliminate the position preference effect. During testing, the mice were deprived of water and food for 12h, followed by the placement of 1% sucrose solution and pure water for another 12h, and the amount of the two solutions consumed in testing phase was measured. The sucrose preference index was calculated as follows: Sucrose preference ratio (%) = Sucrose solution intake (g) / [Sucrose solution intake (g) + Pure water intake (g)] × 100%.

#### OFT

Mice were placed in the center of an open field arena (40×40×40 cm) to freely explore for 10min. The central square area (24×24 cm) was designated as the center zone. The total distance traveled in the open field, entries into the center zone, and time spent in the center zone were automatically tracked using Anymaze software.

#### EPM

The EPM apparatus consists of a cross 50cm above the ground (single side arm 35cm long, 5cm wide, closed arm 15cm high). Mice were placed in the central area with their heads facing the open arm and allowed to freely explore for 5 min. The number of entries into the open arms and the time spent in the open arms were automatically tracked using Anymaze software. The percentage of time spent in the open arms (Open Arm Time Ratio) was calculated as follows: time spent in open arms (s) / [time spent in open arms (s) + time spent in closed arms (s)] × 100%.

#### SIT

The three-chamber test apparatus is an area (40 cm wide, 60 cm long, 20 cm high) divided into three equal areas by two transparent dividers with a shuttle channel at the bottom of the centre. A metal mesh cage is placed in the area on each side of the distal end. The testing process comprised two 10 min phases. During Phase 1 (adaptation phase), the test mouse is placed in the central area with the shuttle passage closed, allowing it to freely explore and familiarize itself with the environment. In Phase 2 (social test phase), an unfamiliar conspecific social mouse is placed in a metal cage on a random side area. The test mouse is then placed in the central area with the shuttle passage open, allowing free exploration of the three areas. Anymaze software automatically tracked the time the mouse spent in each area. The social interaction ratio was calculated as follows: Social interaction ratio (%) = Time spent in the social area (s) / [Time spent in the social area (s) + Time spent in the empty cage area (s)] × 100%.

#### TST

During testing, tape is attached to the tail tip of the experimental mouse, 1 cm from the end. A plastic tube is placed over the mouse’s tail to prevent climbing. The mouse’s tail is then secured to the top of the tail suspension apparatus, suspending the mouse approximately 25 cm above the ground for 6 min. The duration of immobility (s) during the 6 min video recording was recorded and analyzed.

#### FST

The experimental mouse is placed in a transparent cylindrical container (10 cm in diameter, 30 cm high) with water (25 cm deep, 25±1°C) to swim for 6 min. The duration of immobility (s) during the last 4 min of the 6 min video recording was quantified and analyzed.

#### RT

During the training phase, mice were placed on a rotating rod at a constant speed of 5 rpm for 5 min, and the mice were placed back on the rod immediately after falling off the rod to minimize accidental falls from the rod during the test phase. Then mice were rested for 1h. During the test phase, the rotating rod was set to run at an acceleration of 20 rpm/min from 4 rpm to a maximum of 40 rpm for 5 min, and the mice were placed on the rotating rod at the beginning of the initial speed. The formal test consisted of 3 replicate trials with a 30 min rest interval between each trial. The average value of the fall latency (s) of the 3 repeated trials was calculated.

### Molecular biology test

This study employed qRT-PCR and Western blot to assess the mRNA and protein expression levels of GluN3A subunit in different brain regions of mice. Immunofluorescence staining, coupled with confocal imaging, was utilized for staining analysis of neurons in the CA1 region and their projection areas. Additionally, the spatial location of viral expression was confirmed by observing the fluorescence of the injected viruses in each experiment.

#### qRT-PCR

After rapidly decapitating the experimental mice, the brain was promptly dissected and placed in a freezing microtome. Specific brain region boundaries were delineated based on the mouse brain atlas, and target brain regions were extracted using a 1-mm-diameter tissue extraction needle [72], as shown in the Supplementary fig1b. Total RNA was extracted from the tissue samples using TRIzol reagent (Cat# 15596018CN, Invitrogen, USA) following the manufacturer’s instructions, and the concentration was analyzed using NanoDrop (Thermo Scientific, Massachusetts, USA). Subsequently, 500 ng of RNA samples were reverse transcribed using the M-MLV Reverse Transcriptase (Cat# M1701, Promega, USA) following the reverse transcription kit instructions, with a two-step reaction condition: 70°C for 5 min, 37°C for 60 min, to generate cDNA. SYBR Green PCR Master Mix (Cat# CW2601M, Cowin Biotech, China) was used for fluorescence qPCR reactions. The real-time PCR instrument reaction steps were: a single cycle (95°C/10 min); 40 cycles (95°C/15 s; 60°C/1 min); a single cycle (95°C/15 s; 60°C/30 s; 95°C/15 s). Analysis was performed using the ΔΔC(t) method, with GAPDH as the reference gene for standardization. The primer sequences used were: Grin3A: forward: 5’-AGTACCTGAAGAATGATCCAGAGT-3’, reverse: 5’-TCAGATATGTTAGAGGTCAACGG-3’; Gapdh: forward: 5’-GCATGGCCTTCCGTGTTCCT-3’, reverse: 5’-CCCTGTTGCTGTAGCCGTAT-3’.

#### Western blot

The target brain area tissue was homogenized in RIPA lysis buffer containing 1% PMSF (Cat# P0100, Solarbio, China). The collected protein supernatant was quantified using a BCA protein quantification kit (Cat# PC0020, Solarbio, China), and then protein samples were mixed with 4× protein loading buffer (Cat# P1015, Solarbio, China) for denaturation. The equal amounts of samples were loaded onto a 10% SDS-polyacrylamide gel (Cat# CW0022M, Cowin Biotech, China) for separation and then transferred to a 0.2 μm PVDF membrane (Cat# 03010040001, Millipore, Germany). The PVDF membrane was blocked in TBST solution containing 5% milk powder and then incubated overnight at 4°C with diluted primary antibodies (5% milk powder TBST): Grin3A (1:500; Cat# orb694249, Biorbyt, UK); Gapdh (1:3000; Cat# 2118, Cell Signaling, USA). After washing with TBST, the membrane was incubated at room temperature for 2 h with secondary antibodies (5% milk powder TBST) containing horseradish peroxidase: Goat Anti-Rabbit IgG H&L (1:3000; Cat# ZB-2301, Zhongshan Golden Bridge, China). After washing again, the ECL luminescent liquid (Cat# P90719, Millipore, Germany) was added to the PVDF membrane, and the immunoblot signal was captured using an imaging system (FluorChem E, Protein Simple, California, USA).

#### Immunofluorescence staining

90 min after the completion of the FST or CNO injection, mice were intraperitoneally injected with pentobarbital (100 mg/kg) for anesthesia. Following anesthesia, mice were perfused with 30 ml of phosphate-buffered saline (PBS) through the heart, followed by 30 ml of 4% paraformaldehyde (PFA). The brain was removed, fixed in 4% PFA for 24 h at 4°C, and then transferred to PBS with 30% sucrose for dehydration. After the brain settled, 40 μm thick coronal brain slices were obtained using a freezing microtome (CM3050S, Leica, Wetzlar, Germany) and stored at - 80°C in an anti-freezing solution (50% PBS, 30% ethylene glycol, 20% glycerol). For immunofluorescence, the brain slices to be stained were washed with PBS, placed in PBST (0.1% TritonX-100, PBS) containing 5% goat serum at room temperature for 1.5 h, and then transferred to the primary antibody solution for overnight incubation at 4°C. The primary antibodies used were c-Fos (1:500; Cat# SYSY.226003, Synaptic Systems, Germany) and DYKDDDDK Tag (1:500; Cat#14793, Cell Signaling, USA). After washing with PBS, the slices were transferred to the secondary antibody solution at room temperature away from light for 1.5 h. The secondary antibodies used were Goat Anti-Rabbit IgG H&L (Alexa Fluor® 488) (1:500; Cat# ab150077, Abcam, UK) and Goat anti-Rabbit IgG (H+L), Alexa Fluor™ 546 (1:500; Cat#A11010, Invitrogen, USA). After washing with PBS, the slices were incubated in PBS with DAPI (1:500; Cat# C0060, Solarbio, China) at room temperature away from light for 10 min. Following another PBS wash, the slices were mounted on glass slides using a mounting medium (Cat# ab104139, Abcam, UK).

#### Confocal imaging and analyses

Sliced brains or immunofluorescent stained brain slices were imaged using a high-speed confocal imaging system (SpinSR, Olympus, Tokyo, Japan) and a digital slice scanning and analysis system (DMI8, Leica, Wetzlar, Germany) under 10× and 20× objectives. Automated cell counting for c-Fos-positive neurons was performed using the ImageJ Fiji plugin (http://imagej.nih.gov/ij/). Two to four images were selected in each animal for statistical analysis of the brain regions.

### In vivo manipulation

#### AAV

AAV2/9-syn-m-Grin3A-3×flag-Null (titer 1.6×1012 vg/mL) and AAV2/9-syn-ZsGreen (titer 1.6×1012 vg/mL), AAV2/9- syn-hM3D(Gq)-mCherry (titer 1.2×1012 vg/mL), AAV2/9-syn-hM4D(Gi)-mCherry (titer 1.4×1012 vg/mL), and AAV2/9-syn-mCherry (titer 1.5×1012 vg/mL) were purchased from Hanbio Biotechnology, Shanghai, China. AAV2/PHP.eB-mCaMKIIα-GCaMP8s-WPRE-pA (titer 1.1×1013 vg/mL) was purchased from Taitool Bio, Shanghai, China. The viral vectors were aliquoted and stored at -80°C.

#### Stereotaxic surgery

Mice were anesthetized with isoflurane and secured in a stereotaxic apparatus (RWD Life Science, Shenzhen, China). After exposing the skull, the corresponding virus was slowly injected into the target location (AP: -2.90 mm; ML ± 3.30 mm; DV: -2.90 mm) using a Stoelting pump (QSI storage pump, Stoelting, Wisconsin, USA) attached to a Hamilton syringe. After injection, the syringe was left in place for 10 min before slow withdrawal. The mice were allowed to recover for 14 days before behavioral testing.

#### Regulation of GluN3A subunit and virus tracing experiments

### 0.15 µL (unilateral) of AAV2/9-syn-m-Grin3A-3×flag-Null or AAV2/9-syn-ZsGreen was bilaterally injected into CA1i at a rate of 0.05 µL/min

#### Chemogenetic experiments

0.15 µL (unilateral) of AAV2/9-syn- hM3D(Gq)-mCherry or AAV2/9-syn-hM4D(Gi)-mCherry (or AAV2/9-syn- mCherry) was injected into each hemisphere of CA1i at a rate of 0.05 µL/min. 14 days later, CNO (Cat# HY-17366, MCE, USA) was dissolved in dimethyl sulfoxide (DMSO, Cat# D8418, Sigma, Germany) to create a 100 mg/mL stock solution. Before injection, further dilution was made with 0.9% saline to achieve a CNO injection quality of 0.2 mg/kg (injection volume: 10 mL/kg) for chemogenetic activation experiments and 2 mg/kg for chemogenetic inhibition experiments. CNO was intraperitoneally injected 30 min before behavioral testing.

#### Fiber photometry experiments

AAV2/PHP.eB-mCaMKIIα-GCaMP8s-WPRE-pA was diluted 1:4 in cerebrospinal fluid. 0.2 µL (unilateral) of the viral diluted solution was injected into the bilateral CA1i at a rate of 0.05 µL/min. Then, a ceramic optical fiber (200 µm in diameter, 3 mm length, 0.37 NA, Inper Technology Co., Ltd., Hangzhou, China) was implanted above the CA1i region (AP: -2.90 mm; ML ±3.30 mm; DV: -2.85 mm). The fiber was secured to the skull with dental cement. 14 days later, power levels of 470 nm excitation light and 410 nm excitation light in the Inper Signal fiber photometry system (Inper Technology Co., Ltd., Hangzhou, China) were measured using a photometer and adjusted to 25 µW and 10-15 µW, respectively. Prior to the TST, the bundled fiber was connected to the optical fiber implant in the mouse’s head for a two-min adaptation. The Inper Signal system recorded Ca2+ activity in the CA1i region of mice suspended and during tail suspension tests, synchronizing with a camera to record the behavioral state. Inper Plot software was used to analyze and statistically process the fluorescence change data (F) during hanging and struggling events, calculating the relative fluorescence change ΔF/F0 (ΔF/F0 = (F-F0)/F0). Mice with fiber implantation positions confirming the expected target were included in data analysis.

#### Drug delivery experiments

After injecting chemogenetic viruses into bilateral CA1i, 22-gauge single-guide cannulas (RWD Life Science, Shenzhen, China) were bilaterally implanted 0.5 mm above the target positions (AP: -2.90 mm; ML: ±3.30 mm; DV: -2.40 mm). The cannulas were fixed on the skull surface with dental cement. 14 days later, 10 mg/mL D-serine was prepared in CSF. 30 min before the behavioral test, mice were anesthetized with isoflurane, and an injection inner tube exceeding the cannula target position by 0.5 mm was inserted into the mouse’s head cannula. Subsequently, using a dual-channel microinjection pump (RWD Life Science, Shenzhen, China), 0.5 µL (unilateral) of D-serine or CSF was injected into the bilateral CA1i region at a rate of 0.2 µL/min. After completion, the inner tube was left in place for 5 min before slow withdrawal.

### SnRNAseq Analysis

The samples from “Control WT” and “RSD WT” in GSE253687 were reanalyzed using the standard analysis pipeline of Seurat (version 4.4.0)[73]. Briefly, only cells expressing at least 200 genes and genes containing non-zero values as well as associated with at least 3 cells were maintained for subsequent analysis. “NormalizeData” functions were used to normalize data. Highly variable genes were observed using the “FindVariableFeatures” function with default parameters setting, and these highly variable genes were used to perform principal components analysis (PCA). Dimensionality reduction was performed using the “RunUMAP” function, where reduction was set to ‘pca’ and dims to 1:50. Default resolution was used for clustering, using “FindNeighbors” and “FindClusters” functions. Next, we extracted the cells from the CA1 excitatory neuron subpopulation (Mpped1 positive) and performed the aforementioned analysis again. Due to the lack of effective marker genes for screening the CA1 inhibitory neuron subpopulation, we could not extract and analyze the inhibitory neuron subpopulation of CA1[74]. The R package ggplot2[75] was used for visualization.

### Statistical Analysis

All data are presented as mean ± standard error of the mean (M±SEM) and analyzed using GraphPad Prism 9.0 software. Normality distribution was assessed using the Anderson-Darling test. Differences between two groups were evaluated using unpaired two-tailed Student’s t-test, and when variances were unequal, Welch’s correction was applied. Analysis of variance with one or two factors was used for comparisons among three groups or more, followed by FDR: Benjamini-Krieger-Yekutieli correction for multiple comparisons. A *p*-value < 0.05 was considered statistically significant.

## ACKNOWLEDGEMENTS

We are grateful to the lab of Tifei Yuan (Shanghai Mental health center, Shanghai, China) for providing the GluN3A KO mice. We are grateful to Wenyu Tian (Shandong University, Shandong, China) for database technologies and analysis assistance. We greatly appreciate the excellent work of the technical support staff at the Institutional Center for Shared Technologies and Facilities of Institute of Psychology, Chinese Academy of Sciences.

## AUTHOR CONTRIBUTIONS

Methodology, data curation and investigation, writing-original draft&editing: WZ; Research conceptualization, project administration, funding acquisition and editing: JW; Research conceptualization, resources, project administration, funding acquisition, supervision and writing-review&editing: WW; Methodology: MZ, JD, LZ and HY participated.

## ADDITIONAL INFORMATION

## Supplementary information

