## Supplementary information for "Non-classical NMDA receptor subunit GluN3A in a specialized hippocampal region regulates stress-coping strategies in mice"

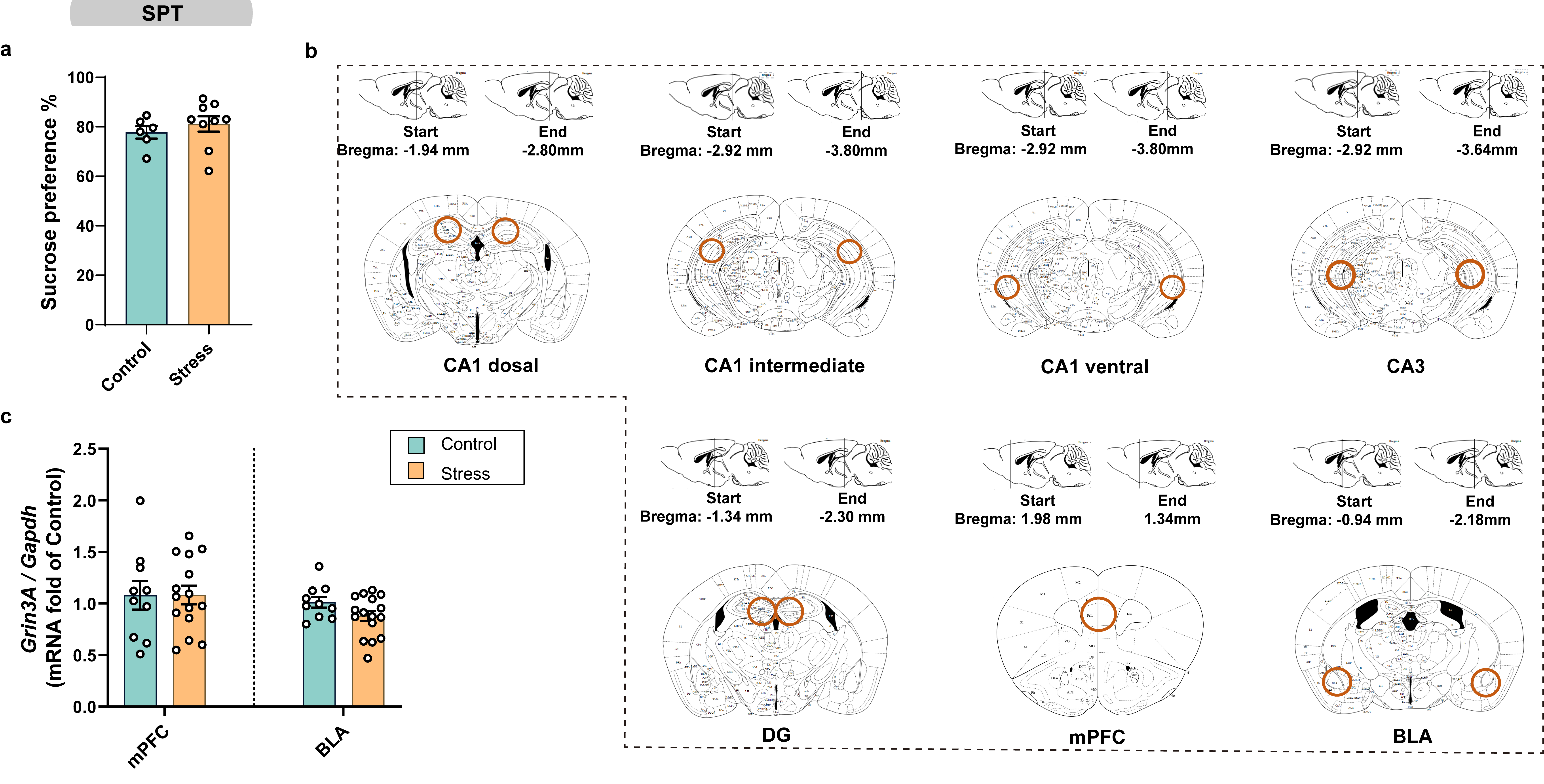

**Supplementary figure 1.** **Chronic social defeat stress (CSDS) did not affect sucrose preference behavior and *Grin3A* mRNA expression level in medial prefrontal cortex (mPFC) and basolateral amygdala (BLA).**

**a** The sucrose preference ratio in sucrose preference test (SPT) (*n =* 6 in Control, *n* = 9 in Stress). **b** Diagram showing dissection of target brain area. Round shape indicating the regions of interest in brain slices. **c** Quantitative data of *Grin3A* mRNA in target brain area from control and stress mice (mPFC: *n =* 10 in Control, *n* = 15 in Stress; BLA: *n =* 10 in Control, *n* = 16 in Stress).

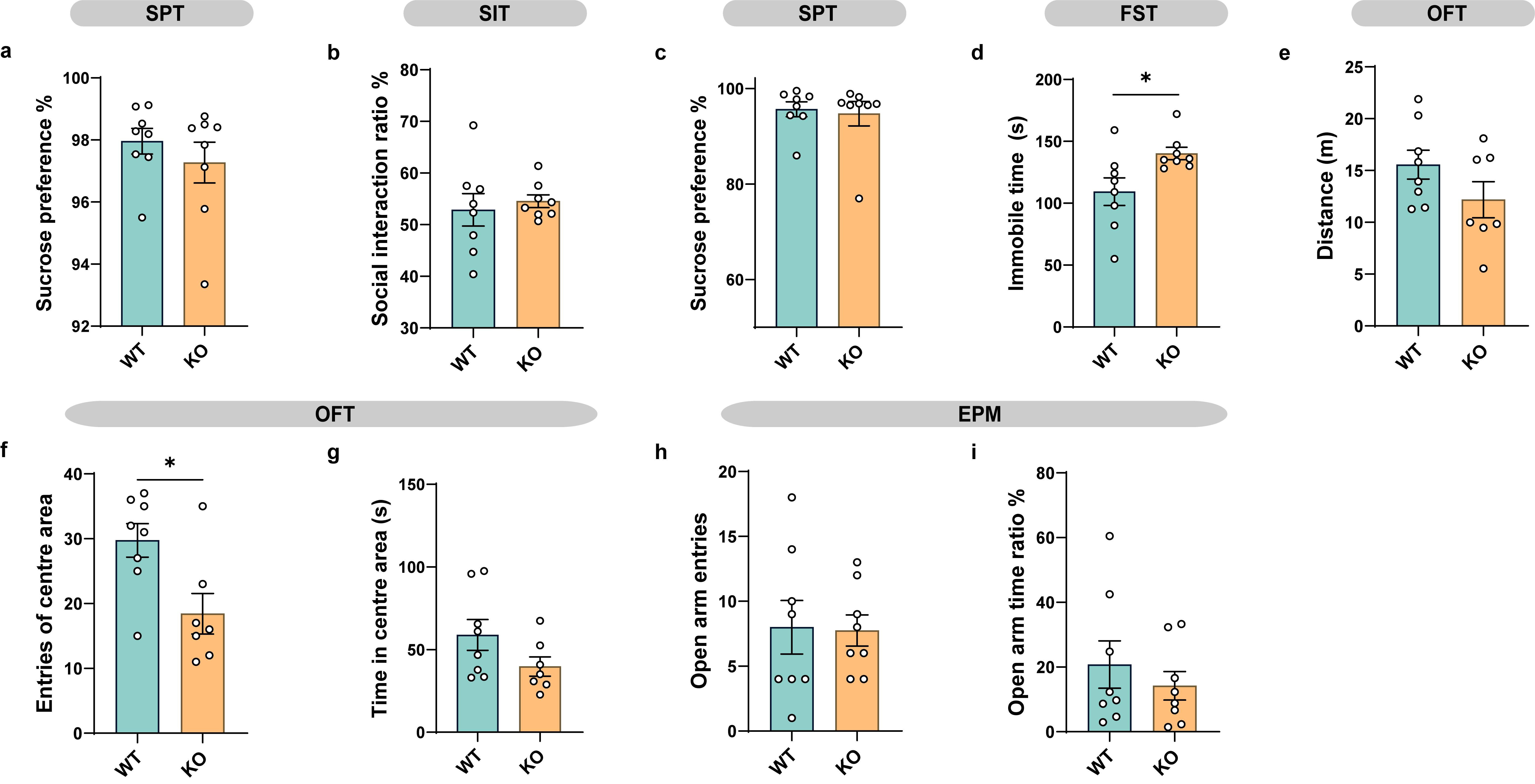

**Supplementary figure 2. GluN3A knockout (KO) mice, whether experiencing subthreshold social defeat stress (SDS) or not, showed no alteration in emotion-related behaviors except for coping behaviors.**

**a** The sucrose preference ratio in SPT (*n* = 8 per group). **b** Social interaction ratio of social interaction test (SIT) (*n* = 8 per group). **c** The sucrose preference ratio in SPT (*n* = 8 per group). **d** Immobility time in forced swim test (FST) (*n* = 8 per group). **e** The total distance traveled in open field test (OFT) (*n =* 8 in WT, *n* = 7 in KO). **f** The number of entries to centre area of OFT. **g** The time in the centre area of OFT. **h** The number of entries to open arms of elevated plus maze (EPM) (*n* = 8 per group). **i** Percentage of time spent in the open arms of EPM. **P* < 0.05.

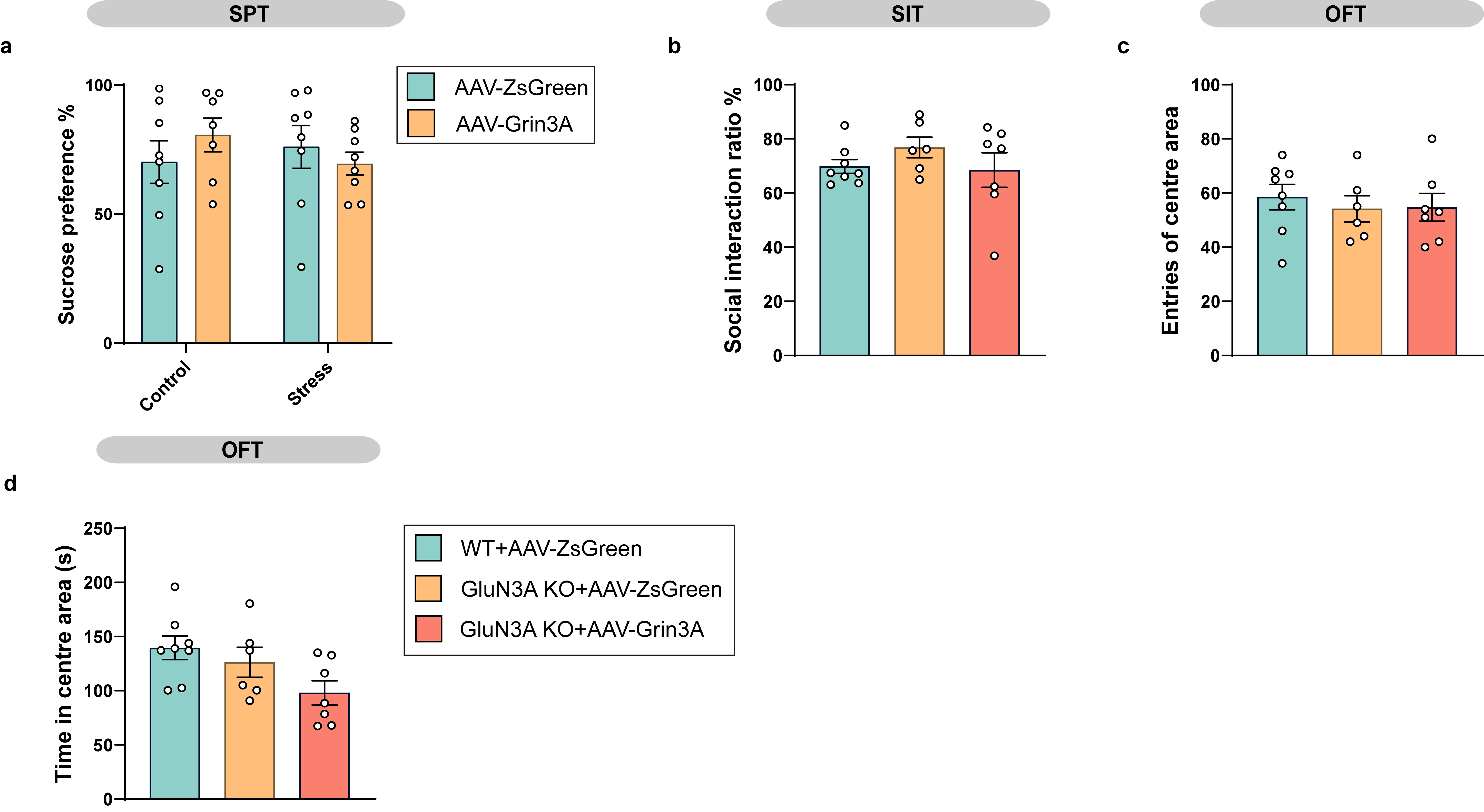

**Supplementary figure 3. Overexpression of GluN3A in CA1 intermediate (CA1i) did not affect other emotion-related behaviors.**

**a** The sucrose preference ratio in SPT (*n* = 8, 7, 8 and 8 in control:AAV-ZsGreen, control:AAV- Grin3A, stress:AAV-ZsGreen and stress:AAV-Grin3A, respectively). **b** Social interaction ratio of SIT (*n* = 8, 6 and 7 in WT+AAV-ZsGreen, GluN3A KO+AAV-ZsGreen and GluN3A KO+AAV- Grin3A, respectively). **c** The entries of centre area in OFT (*n* = 8, 6 and 7 in WT+AAV-ZsGreen, GluN3A KO+AAV-ZsGreen and GluN3A KO+AAV-Grin3A, respectively). **d** The time in the centre area of OFT.

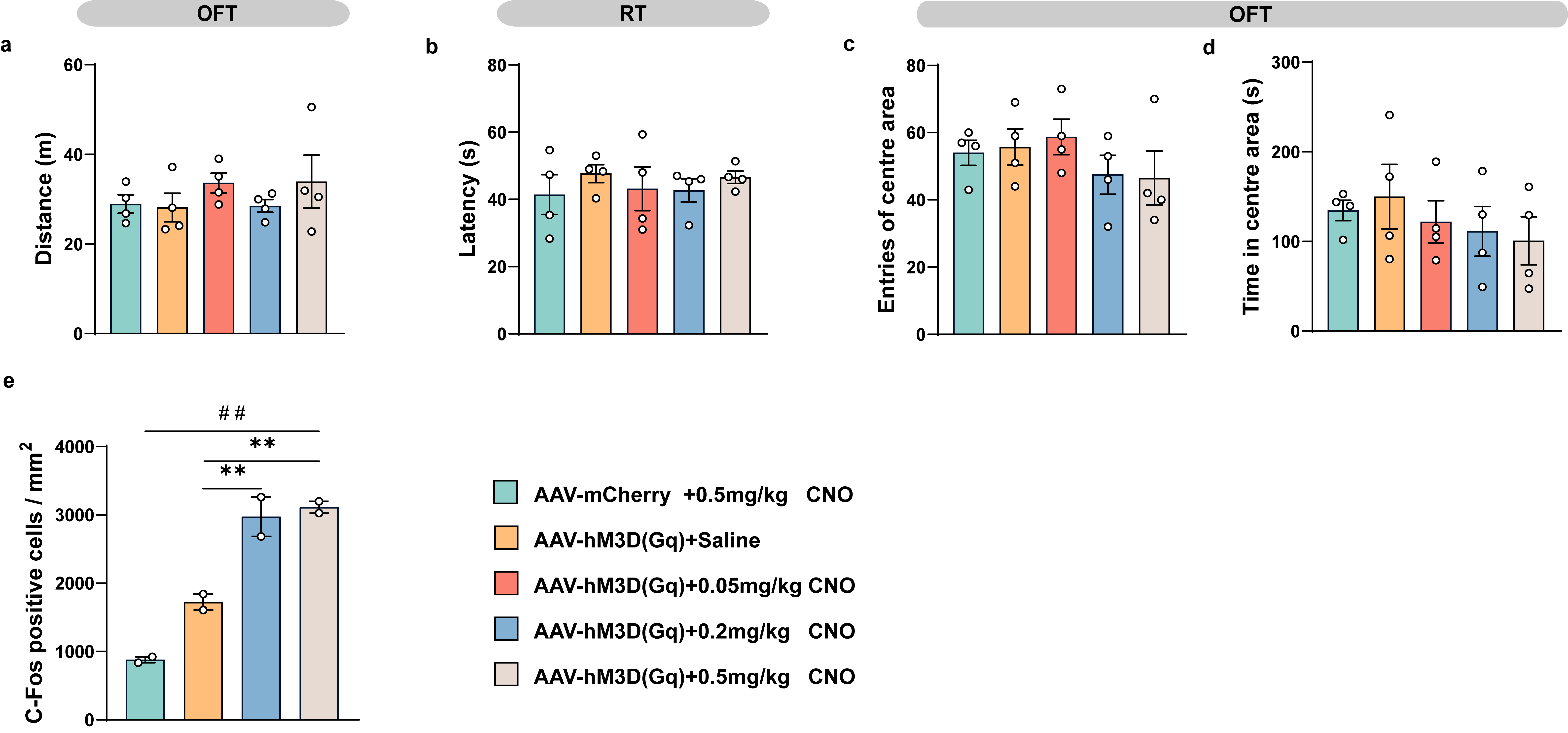

**Supplementary figure 4. Clozapine N-oxide (CNO) concentration gradients effect of chemogenetic activated virus on mice locomotor activity and c-Fos-positive cells in CA1i pyramidal layer neurons.**

**a** The total distance traveled in OFT (*n* = 4 per group). **b** the latency of falls in rotarod test (RT) (*n* = 4 per group). **c** The entries of centre area in OFT. **d** The time in the centre area of OFT. **e** Quantification of c-Fos-positive cells in CA1i pyramidal layer neurons of each group of mice. (*n* = 2 per group, 4 slices for each dot). *** P* < 0.01, *^##^ P* < 0.01.

### **Supplementary figure 5**. GluN3A-mediated activation of CA1i pyramidal neurons **did not** influences neuronal activity in downstream nucleus accumbens (NAc) circuit.

**a** Statistical region of c-Fos-positive cells in NAc core (AcbC) and shell (AcbSh) (ZsGreen: green, c-Fos: red, DAPI: blue). Scale bar, 500 μm. **b** Quantification of c-Fos-positive cells in AcbSh neurons of each group of mice (n = 4, 5, 5 and 4 in control:AAV-ZsGreen, control:AAV-Grin3A, stress:AAV-ZsGreen and stress:AAV-Grin3A, respectively, 4 slices for each dot). **c** Quantification of c-Fos-positive cells in AcbC neurons of each group of mice (n = 4, 5, 5 and 4 in control:AAV-ZsGreen, control:AAV-Grin3A, stress:AAV-ZsGreen and stress:AAV-Grin3A, respectively, 4 slices for each dot). **d** Representative images of c-Fos-positive cells in the AcbSh and AcbC of each group of mice (c-Fos: red, Dapi: blue). Scale bar, 50 μm.
